# Physical activity promotes gut adaptation, responses to nutrients, and sensitivity to gut peptides

**DOI:** 10.1101/2025.05.12.653520

**Authors:** Cecilie Bæch-Laursen, Jon Vergara Ucin, Katrine Douglas Galsgaard, Jesus Llana, HanneLouise Kissow, Jens Juul Holst, Bente Klarlund Pedersen, Paula Sanchis

## Abstract

Physical activity is essential for body weight maintenance after body weight-loss, partly by promoting the coupling between energy intake and expenditure. However, the underlying mechanisms remain largely unknown. Here we demonstrate that running induces small intestine growth independently of GLP-2. In addition, exercise increases L-cell density in the small intestine and glucose-stimulated GLP-1 secretion, and improves the sensitivity both to the gut-derived hormones PYY, CCK and ghrelin, and to treatment with GLP-1 receptor agonist. Moreover, increased physical activity enhances satiation and satiety post-fasting, regulates the gene expression of appetite signals in the intestine, nodose ganglia and brainstem, and induces a greater feeding-response in the activation of hypothalamic and brainstem neurons. This improves overall appetite regulation, and in turn, promotes body weight maintenance. In summary, the present data suggest that increased physical activity improves body weight maintenance by inducing adaptations in the gut and in gut-to-brain communication that control appetite.

**HIGHLIGHTS:**

1. Increased physical activity promotes intestinal growth
2. Sensitivity to gut-derived hormones PYY, CCK and ghrelin is improved in active mice
3. Increased physical activity enhances glucose-stimulated GLP-1 secretion from the small intestine
4. Increased physical activity regulates effectively satiation and satiety post-fasting

## Introduction

Following an initial diet-induced body weight-loss, ∼70% of the lost weight is typically regained within five years emphasizing the significant challenge of long-term weight maintenance.^1^ Retrospective studies have demonstrated that physical activity is associated with a greater chance of maintaining weight-loss and thus preventing body weight gain.^2^ While the acute calorie deficit from a single exercise bout may not alone be sufficient for body weight maintenance,^3–5^ increasing the level of daily physical activity may influence appetite regulation and promote a more accurate coupling of energy intake and expenditure.^6^ Indeed, energy intake seems to have a J-shaped relationship with physical activity levels, so that low levels of physical activity result in dysregulated appetite and high regular levels associate with better appetite control.^7^ Thus, exercise does not only increase food intake because of higher energy expenditure but also regulates subjective hunger feelings, enhances post-meal satiety, and reduces the overconsumption of calorie-dense food.^8,9^

Significant progress has been made in understanding how nutrient-stimulated peripheral signals are relayed to the brain to control feeding behavior and thus body weight maintenance.^10^ The gastrointestinal tract, from where several satiety- and hunger-promoting signals derive, is central for sensitive appetite regulation. Among these hormones, glucagon-like peptide-1 (GLP-1), peptide YY (PYY) and cholecystokinin (CCK) are secreted in response to a meal and induce satiety.^11–16^ GLP-1 is co-secreted with PYY from L-cells of the small intestine and colon, and CCK is secreted from the upper small intestine. The signal from the gut to the brain can be transmitted either through the gut hormones release to bloodstream or through the afferent part of the vagus nerve. Whereas endogenous GLP-1 and CCK act mainly on vagal afferents to reduce food intake, PYY impacts directly on the brainstem dorsal vagal complex (DCV), mainly in the area postrema (AP), and on the hypothalamic *Agrp*- and *Pomc*-expressing neurons, and the activity hereof are determining food intake.^17–19^ Ghrelin, another nutrient-stimulated peripheral hormone secreted from the stomach, the circulating levels increased before meals enhancing appetite by impacting to the hypothalamus and DCV, in part in nucleus of the solitary tract (NTS) and AP.^19^

It seems that acute and chronic exercise differentially modulates gut-derived hormones. Acute exercise increases circulating levels of PYY and GLP-1 in fed-state healthy humans,^8^ whereas habitual physical activity decreases circulating levels of GLP-1 and enhanced glucose-stimulated GLP-1 secretion, independent of insulin sensitivity, in fasted-overweight men.^20^ CCK increases with acute exercise in healthy males but was not modified by repeated bouts of exercise, however the evidence is based on a limited number of studies.^21,22^ Ghrelin is generally increased following long-term chronic exercise and decreased after high-intensity acute exercise.^23,24^ Importantly, exercise combined with pharmacological treatment with GLP-1 receptor agonists (GLP-1RA),^25^ effectively promotes additional weight-loss maintenance and potentiates fat-loss compared to groups treated with either GLP-1RA or one year exercise alone.^26^ These changes may constitute the link between physical activity and energy intake. Thus, it is possible that the increased energy intake resulting from activity-induced higher expenditure drives gastrointestinal adaptations that modulate these nutrient-stimulated peripheral hormones and coordinate communication with the brain, ultimately ensuring body weight maintenance.

In the present study, we have uncovered that voluntary wheel running induces adaptations in the gut and in gut-to-brain communication in mice. After 6 weeks of running, active mice displayed higher food intake, positive changes on body composition without an overall effect on body weight, and a better matching of energy intake to expenditure in *ad-libitum* fed conditions and following fasting intervention. Active mice displayed longer small intestine and colon, along with greater sensitivity to the gut-derived hormones PYY, CCK, and ghrelin despite lower levels of circulating PYY and ghrelin were observed with running. Interestingly, active mice had a greater increase in glucose-stimulated GLP-1 secretion and higher density of GLP-1 expressing L-cells. Moreover, increased physical activity improved satiation and satiety responses after calorie restriction and modulated the molecular landscape along small intestine, colon, nodose ganglia and brainstem, being the last one also regulated by feeding state. Acutely, a greater feeding-response in the activation of hypothalamic and brainstem neurons was also observed due to increased physical activity. Altogether, our data demonstrate that increased physical activity induces gut adaptations which in turn influences appetite regulation and body weight maintenance.

## Results

### Physical activity induces small intestine and colon growth

First, we assessed the physiological response to regular physical activity by exposing male mice to 6 weeks of voluntary wheel running. A*d-libitum*-fed mice were single-housed, and based on equal phenotypic distribution, they were assigned to active (functional running wheel) or sedentary (blocked running wheel) groups (Figure 1A). Active mice ran on average 257 km over a 6-week period corresponding to 6.1 km/day and the running distance correlated positively with food intake, increasing by 0.43 g per km (Figure 1B). Active mice consumed 0.721±0.1 g/day, in relative terms 20% more food compared to sedentary controls (Figure 1C). Body weight increased by 1.18±0.25 g in sedentary mice and by 1.55±0.25 g in active mice during the 6 weeks (Figure 1D). After 6-weeks of wheel running, active mice had increased their lean mass by 1.33±0.2 g and decreased fat mass by 0.52±0.2 g whereas sedentary mice increased their lean mass by 0.34±0.1 g and fat mass by 0.30±0.2 g (Figure 1E). Together, these results highlight that voluntary wheel running promotes food intake and improves body composition without overall changes in body weight.

**Figure 1.**
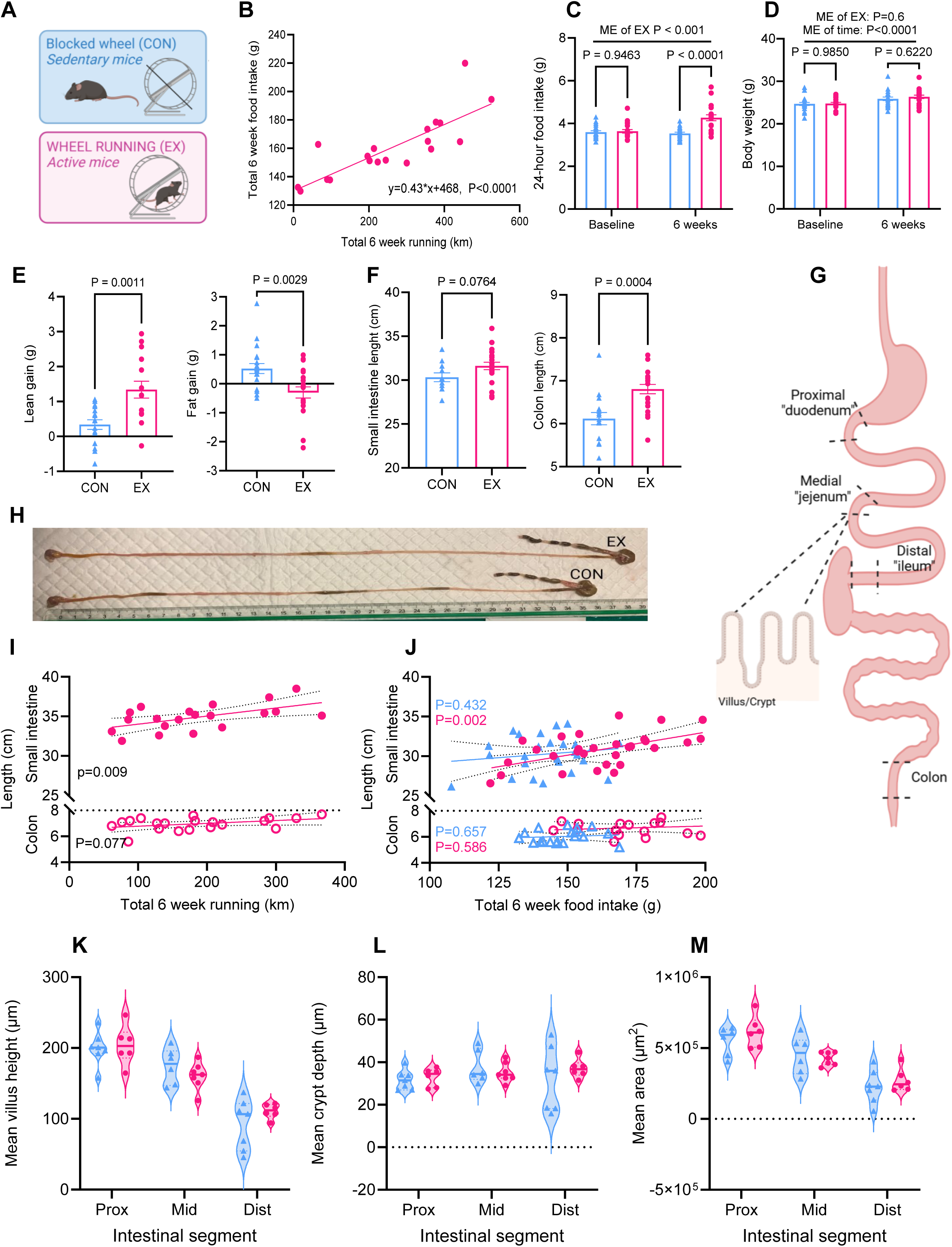
Food intake, body mass, body composition and intestinal length. (A) Color scheme; sedentary mice (mice with a blocked running wheel) are shown in blue and active mice (voluntary wheel running) are shown in pink. (B) Simple linear regression analysis of total 6-week food intake (g) in relation to total 6-week running distance (km). (C) Mean 24-hour food intake in sedentary controls and active mice at baseline and after 6 weeks of running (D) Mean body weight (g) at baseline and at 6 weeks of wheel running or blocked wheel compared by mixed model for repeated measurements followed by Bonferroni multiple comparison. (E) Lean mass gain (g) and fat mass gain (g) after 6 weeks intervention compared by student t-test. (F) Small intestine and colon length in *ad-libitum* fed mice after 6-week intervention compared by student t-test. (G) Illustration of the intestinal segments analyzed. (H) Small intestine with cecum and colon attached from an active and a sedentary mouse. (I) Simple linear regression analysis of the relationship between intestinal length; small intestine (bold symbols) and colon (open symbol) and total 6-week running distance, and (J) the correlation between intestinal length and total 6-week food intake. (K) Villus height µm, (L) crypt depth µm, and (M) luminal area µm^2^ in the proximal, middle, and distal small intestine. Data are shown as mean±SEM. P-values are indicated on the individual graphs. ME of EX is main effect of wheel running.

Given the daily increase in food intake in the running mice, we next assessed whether physical activity induces adaptations in the gastrointestinal tissue. Small intestine was 1.3±0.71 cm longer in *ad-libitum*-fed active mice compared to *ad-libitum*-fed sedentary (P=0.076 for the difference), and the colon was 0.57±0.26 cm longer in active mice (Figure 1F-H). Interestingly, the small intestine length was positively correlated with total 6-week running distance and food intake (Figure 1I-J). There was no correlation between small intestinal length and total 6-week food intake in sedentary mice (Figure 1J), revealing that increased physical activity promotes intestinal growth driven by increased food intake. Histological analysis showed that running did not affect overall or section-specific villus height, crypt depth, or luminal area (Figure 1K-M). In the colon, while the length was positively correlated with total running distance (Figure 1I), there was no correlation between food intake and colon length in either active or sedentary mice (Figure 1J). Overall voluntary wheel running induced intestinal adaptations in mice.

### Running-induced intestinal growth is independent of GLP-2

Glucagon-like peptide-2 (GLP-2) stimulates intestine growth.^27^ Therefore, we evaluated whether GLP-2R deletion affects the training-induced adaptations by exposing male whole-body GLP-2R knockout mice (*Glp2r^−/-^* mice) and their littermate (*Glp2r^+/+^* mice) to 6 weeks of voluntary running (Figure 2A). GLP-2R deletion did not impact on accumulative running distance (Figure 2B), food intake (Figure 2C-D) or body weight (Figure 2E) in sedentary nor active mice, suggesting that the physiological response to running is not influenced by GLP-2. There was no difference in small intestine length between the genotypes in either sedentary or active conditions. The small intestine length increased equally with exercise in *Glp2r^+/+^* (difference of means: 2.02±0.9 cm, P=0.18) and in *Glp2r^−/-^* (difference of means: 1.83±0.87 P=0.22; Figure 2F). Surprisingly, colon length was not affected by genotype nor running, in contrast to the results presented above (Figure 2G), likely resulting from differences in the running distance and smaller sample size. Correlation of colon length to total running distance revealed an effect of running in *Glp2r^−/-^,* and a tendency of a positive effect of running in *Glp2r^+/+^* mice (Figure 2H). Together, we conclude that training-induced lengthening of the small intestine is independent of *Glp-2r* expression.

**Figure 2.**
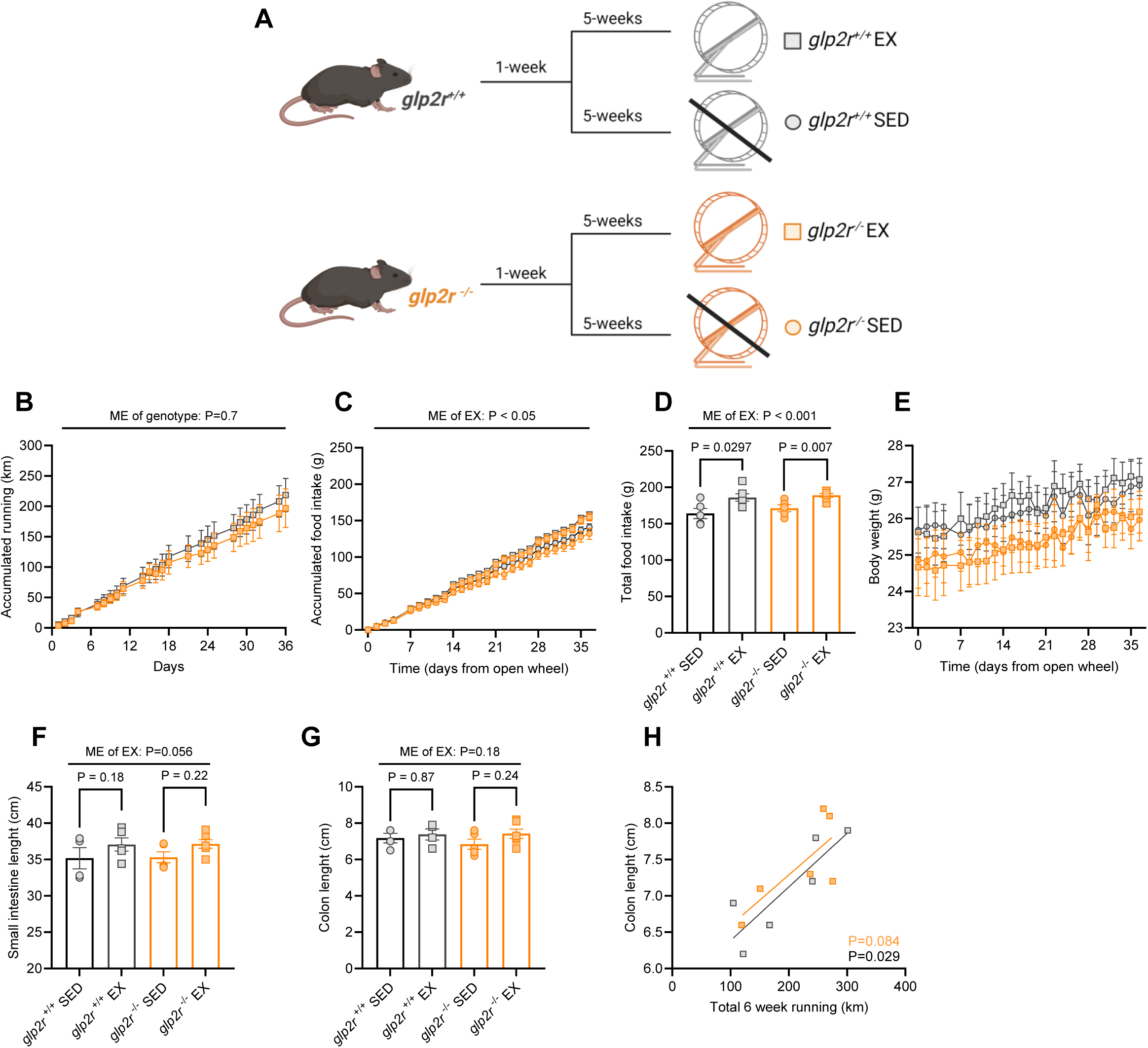
GLP-2 is not essential for exercise induced intestinal lengthening. (A) Schematic overview of the study setup and group colors. Day 0 indicates the day when the wheels were opened for the active mice. (B) Accumulated running distance (km) for 36 days, and (C) accumulated food intake (g) for 36 days compared by mixed-model for repeated measurements. (D) Total food intake at day 36 compared by mixed model for repeated measurements with Bonferroni multiple comparison. (E) Body weight (g) for 36 days. (F) Small intestine and (G) colon length compared by two-way ANOVA. (H) Correlation of total running distance and colon length with simple linear regression model. Data are shown as mean±SEM. P-values are indicated on the individual graphs. ME of EX is main effect of wheel running.

### Increased physical activity promotes the intestinal adaptive response to nutrients

Since voluntary running induces gut adaptations, we then analyzed how nutrient-stimulated peripheral signals are regulated in active mice under *ad-libitum* fed conditions before dark phase. Sedentary and active mice showed similar levels of circulating GLP-1, however circulating levels of PYY and ghrelin were lower in active mice compared to sedentary (Figure 3B-C). Secondly, we investigated glucose-stimulated GLP-1 and PYY secretion through *in situ* isolated small intestine perfusion (Figure 3D and S1A-B). GLP-1 secretion increased in response to a 15-min luminal infusion of glucose in both sedentary and active mice. Glucose-stimulated GLP-1 secretion was 179.1±53.4 pmol higher in active compared to sedentary mice in the same 15-min period (Figure 3E). The greater GLP-1 secretion was independent of small intestine length (Figure S1C). Glucose did not promote PYY secretion neither in active nor sedentary mice (Figure S1A-B). Thirdly, we investigated the number of L-cells by immunostaining for GLP-1 positive cells in the small intestine given their role in communicating satiety (Figure 3F). Density of GLP-1 positive L-cells increased along the small intestine, reaching a peak in the distal ileum, as previously described.^28^ Active mice seemingly had a greater number of GLP-1 positive cells compared to sedentary mice, which was significantly greater in the distal small intestine (Figure 3F-G). In summary, these results emphasize an improved capacity to modulate nutrient-stimulated peripheral signals with increased physical activity.

**Figure 3.**
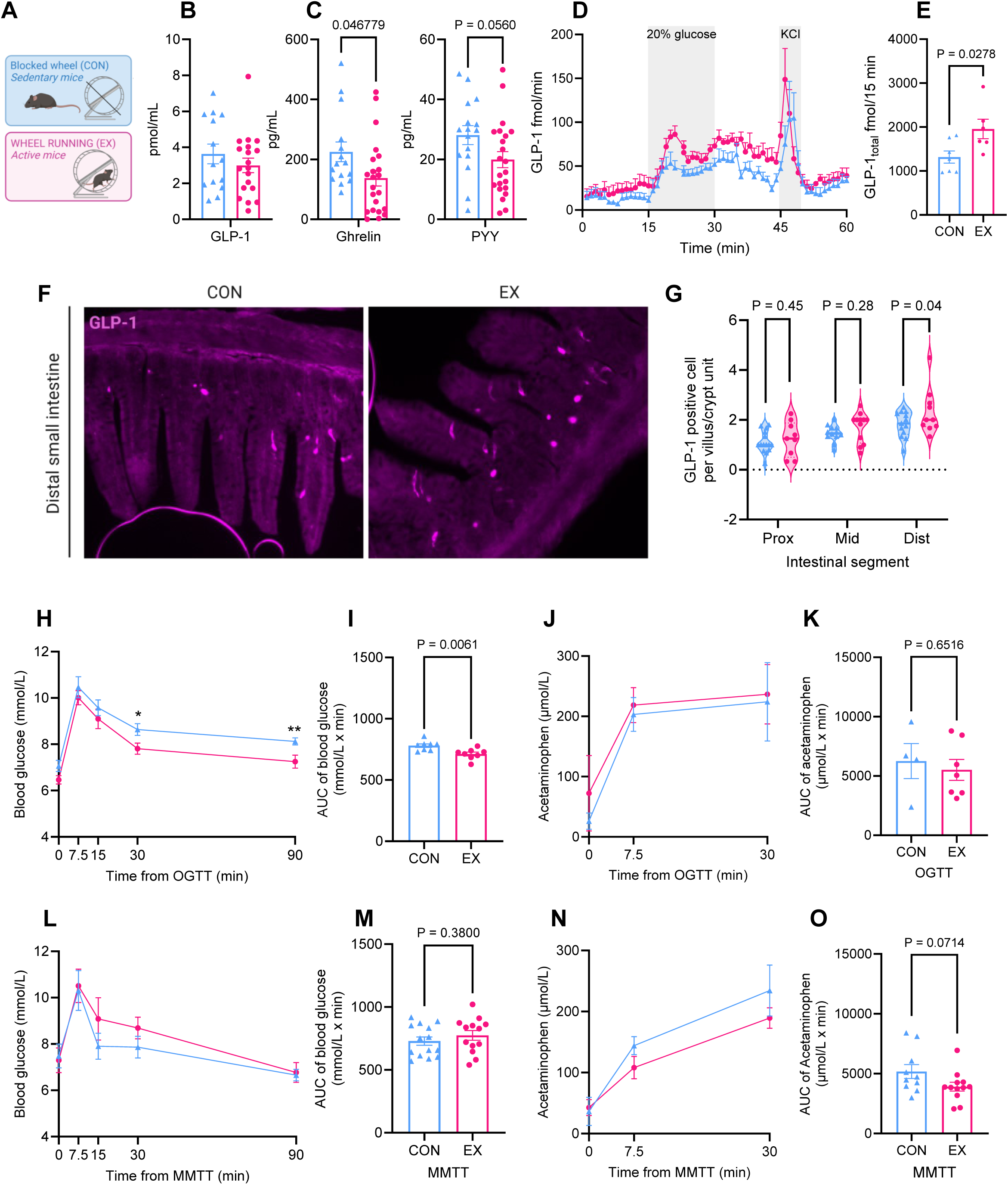
Basal and glucose induced concentration of gut hormones and meal tolerability. (A) Color scheme. (B) *Ad-libitum* feeding serum levels of GLP-1 (pmol/mL), (C) ghrelin and PYY (pg/mL) in sedentary and active mice compared by student t-test. (D) GLP-1 secretion (pmol/L/min) from *in situ* isolated perfused small intestine stimulated with 20% luminal glucose and positive control KCl, (E) total GLP-1 secretion (pmol/15 min) during 15-min luminal glucose stimulation compared by t-test. (F) Representative picture of GLP-1 staining in the distal small intestine of a sedentary and an active mouse. (G) Quantification of GLP-1 positive cells per villus/crypt unit compared by t-test for each segment (CON *n*=6, EX *n*=5). (H) Blood glucose in sedentary and active mice following OGTT. (I) AUC of blood glucose following OGTT compared by t-test. (J) Acetaminophen concentration in serum following OGTT and (K) AUC of acetaminophen absorption following OGTT compared by t-test. (L) Blood glucose in sedentary and active mice following a liquid MMTT. (M) AUC of blood glucose following a MMTT compared by t-test. (N) Acetaminophen serum concentrations following MMTT, and (O) AUC of acetaminophen absorption following MMTT compared by t-test. Data are shown as mean±SEM. P-values are depicted on the graphs.

GLP-1 suppresses food intake and slows gastric emptying.^29^ Given the enhanced glucose-stimulated GLP-1 secretion in active mice, we next examined whether running influences glucose tolerance and gastric emptying. We assessed this using an oral glucose tolerance test (OGTT) and a liquid mixed meal tolerance test (MMTT) combined with acetaminophen as an indicator of gastric emptying rates. Active mice had significantly better glucose tolerance following an OGTT (Figure 3H-I), however there was no apparent effect of running on gastric emptying rates in the OGTT (Figure 3J-K). Glucose tolerance appeared similar between groups during the MMTT (Figure 3L-M). Nonetheless, there was a slight delay (P=0.071 for the difference) in acetaminophen absorption rates in active mice compared to sedentary (Figure 3N-O). The results suggest that running may affect glucose tolerance and perhaps gastric emptying depending on the gastric load employed.

### Increased physical activity enhances sensitivity to gut-derived pharmacological peptides

Given the changes in self-reported feeling of hunger and satiety humans in response to exercise^6,30^ and here showing changes in circulating levels of nutrient-stimulated gut-derived peptides in active mice, we speculated that increased physical activity may induce an enhanced sensitivity to gut-derived hormones. Thus, we conducted a crossover design study in which we monitored food intake following PYY, CCK, and ghrelin injections at the onset of hunger (ZT12) and fullness (ZT0) in *ad-libitum*-fed active and sedentary mice (Figure 4A). Preliminary experiments in sedentary mice were conducted with PYY and CKK, and corresponding vehicles solutions, and Ghrelin was tested in two concentrations in active and sedentary mice to determine a suitable dose of each peptide, which tended to suppress/increase food intake (Figure S1D-F).

**Figure 4.**
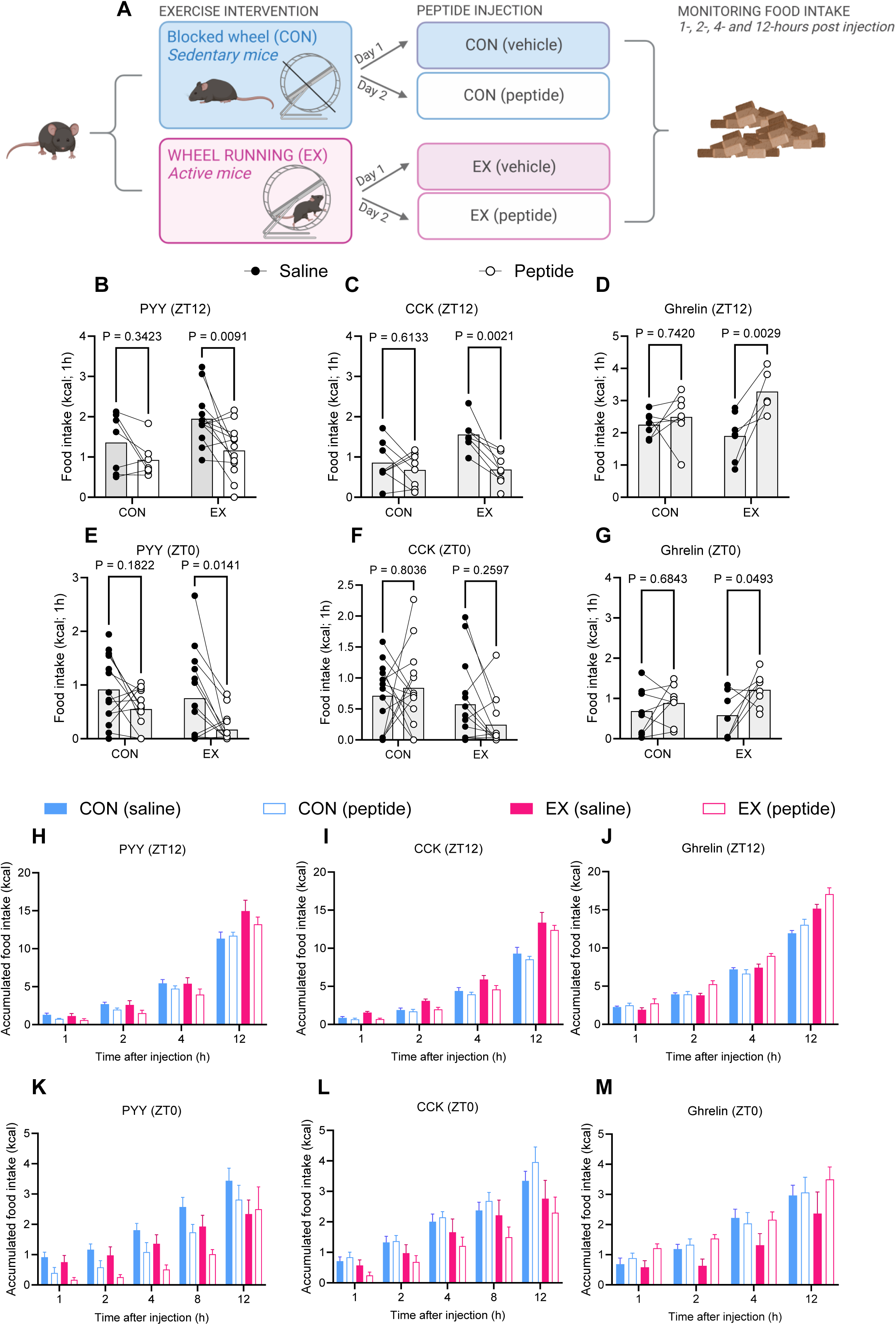
Peptide hormone sensitivity. (A) Schematic overview of experimental setup. (B) 1-hour food intake post saline or PYY injection at ZT12, (B) 1-hour food intake post saline or CCK injection at ZT12, (D) 1-hour food intake post saline or ghrelin injection at ZT12. (E) 1-hour food intake post saline or PYY injection at ZT0, (F) 1-hour food intake post saline or CCK injection at ZT0, (G) 1-hour food intake post saline or Ghrelin injection at ZT0. 1-hour food intake following injection was compared to food intake following vehicle injection by paired t-test. Food intake was measured for 12 hours post injection. (H) Accumulated food intake for 12 hours post PYY injection at ZT12, (I) accumulated food intake for 12 hours post CCK injection at ZT12, (J) accumulated food intake post for 12 hours Ghrelin injection at ZT12. (K) Accumulated food intake for 12 hours post PYY injection at ZT0, (L) accumulated food intake for 12 hours post CCK injection at ZT0, (M) accumulated food intake for 12 hours post Ghrelin injection at ZT0. Data are shown as mean±SEM. P-values are depicted on the graphs.

In the first hour of feeding post-injection at ZT12, PYY suppressed food intake by 40% (Figure 4B), CCK suppressed food intake by 55% (Figure 4C), and Ghrelin increased food intake by 70% (Figure 4D) in active mice. In the first hour of feeding post-injection at ZT0, PYY suppressed food intake by 75% (Figure 4E), CCK trended to suppress food intake by 55% (Figure 4F) and Ghrelin increased food intake by 110% (Figure 4G) in active mice. Importantly, the effects of peptide injection persisted throughout the injection day in active mice, whereas neither of the peptides affected food intake in sedentary mice acutely or in the 12-hours post injection (Figure 4H-M). We were unable to detect suppressed food intake after native GLP-1 injection in either group (Figure S1G), supposedly due to short half-life of GLP-1.^31–33^ Active mice generally ate more during the dark phase and less during the light phase, which reinforces the idea that physical activity enhances hunger signals during the active phase and, conversely, increases satiety during the resting phase. Although administered in similar doses; PYY, CCK and ghrelin had greater effects on food intake in active mice, with the effects being more pronounced during the light phase than the dark phase one-hour post-injection. Together, our data emphasize that increased physical activity enhances sensitivity to anorectic and orexigenic gut-derived hormones.

As described above, gut signaling may influence neural activity of the hypothalamus and brainstem DCV.^17,18^ In particular, PYY elicits its effect on hypothalamic Arcuate nucleus (ARC), mainly in the NPY-receptor positive neurons to suppress food intake, whereas ghrelin acts on hypothalamic AgRP neurons to increase food intake.^19,24,34^ Both PYY and ghrelin have actions on the brainstem, PYY mainly in NTS and ghrelin in NTS and AP.^35,36^ Thus, we evaluated the acute neuronal activity in the ARC and brainstem one-hour post injection of PYY, ghrelin, or their vehicle in active and sedentary mice. Acute neuronal activity was assessed by counting the number of cFOS-positive cells in the relevant brain regions (Figure S1H-J).^18,19^ cFOS activity following PYY or ghrelin injection was compared to the same vehicle-group, as all injections were performed simultaneously. PYY injection resulted in acute neuronal activity in the NTS, and AP, and ghrelin injection resulted in an acute spike in neuronal activity in ARC, AP and NTS. There were no clear effects of running on neuronal activity in the ARC nor NTS, though there was a small non-significant effect in the AP (Figure S1H-J), however it is possibly driven by low sample size in the saline group. In the small intestine, we analyzed expression of *Gcg, Pyy,* and *Cck* following saline injection. *Gcg* and *Cck* expression was greatly upregulated in the proximal and middle small intestine (Figure S1K-M). In conclusion, active mice are more sensitive to the gut-derived hormones PYY, Ghrelin, and CCK, with this sensitivity being likely unrelated to acute neuronal changes.

### Increased physical activity improves satiation and satiety responses after calorie restriction

Increased physical activity improves the sensitivity of anorectic and orexigenic gut-derived hormones during *ad-libitum* feeding (Figure 4). Thus, we hypothesized that running could improve the satiation and satiety during the refeeding after fasting period. This paradign^37^ allowed us to explore the effect of running in the transition from a state of negative energy balance to acute energy intake.^34^ Mice were fasted for 12-hours (ZT0–ZT12) during the light phase and then fed in *ad-libitum* conditions. Food intake and body weight were monitored in the following days and compared to food intake in the 24 hours pre-fasting (Figure 5A-B). The rationale for removing the food during the light-cycle but not dark-cycle was to avoid a strong fasting signal that may mask the differences between sedentary and active mice. After 12-h daytime fasting, the sedentary mice compensated for the energy lost by increasing their food intake by 46% in the following 24-hours, whereas the active mice showed only a small and non-significant increase of 7%. The sedentary mice continued to overconsume the following days, by 25% at 48-hours and by 12% at 72-hours post-fasting, whereas no changes in food intake were observed in active mice over days (Figure 5B). As expected, the compensatory eating response in sedentary mice resulted in additional body weight gain (Figure 5C). Together, these results reveal that satiation and satiety responses are stronger regulated in active mice, even during negative energy-balance, which helps prevent body weight rebound.

**Figure 5.**
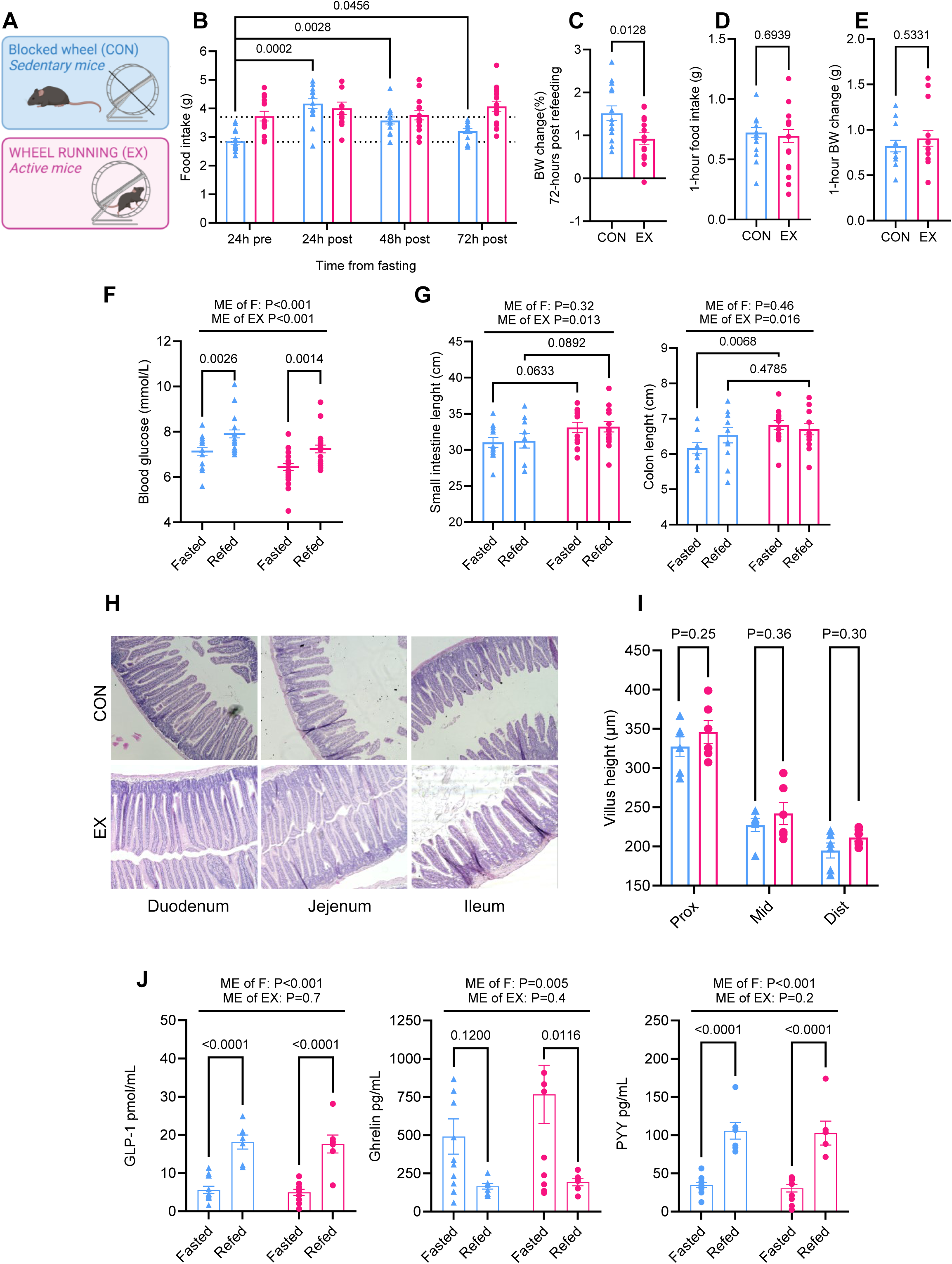
Post-fasting food intake and effects on blood glucose and gut hormones. (A) Color scheme for the figure, showing sedentary mice in blue and active mice in pink. (B) The difference in food intake between 24-hours before and 24-, 48, and 72-hours post a 12-hours daytime fast was compared by Holm-Sídak multiple comparison. (C) Body weight change (%) at 72-hours post 12-hour daytime fast compared by t-test. In a new cohort we investigated fasting and acute 1-hour refeeding. (D) 1-hour food intake, and (E) body weight change post 1-hour refeeding following a 12-hour daytime fast compared by t-test. (F) Blood glucose during fasting and after 1-hour refeeding compared by two-way ANOVA with Bonferroni multiple comparison. (G) Small intestine and colon length in fasted and 1-hour refed mice compared by two-way ANOVA with Bonferroni multiple comparison. (H) Hematoxylin and eosin staining of intestinal tissue. (I) Villus height in 12-hour daytime fasted mice was compared by t-test for each segment. (J) Serum concentrations of GLP-1, ghrelin and PYY in fasted and 1-hour refed mice compared by two-way ANOVA with Bonferroni multiple comparison. Data are shown as mean±SEM. P-values are depicted on the graphs. ME of EX is main effect of wheel running, ME of F is main effect of refeeding.

To evaluate whether the coupling of food intake is improved in other contexts of negative energy balance, we conducted an experiment administering the GLP-1RA semaglutide^38^ to sedentary and active mice (Figure S2A). Semaglutide induced a strong suppression of food intake for 48 hours in both active and sedentary mice (Figure S2B). However, the feeding suppression tended to be stronger in active mice and the body weight-loss to be greater in active mice at 48 hours post injection (Figure S2C), indicating that active mice are more sensitive to the peptide. Indeed, exercise has already been described to potentiate GLP-1RA effects on body weight maintenance.^26^ After 3 days, the baseline food intake was restored in both groups, but whereas sedentary mice were overconsuming by 38%, active mice did not significantly increase their food intake (12%, P=0.8) compared to baseline food intake. Together, the data suggest that not only are GLP-1RA effects greater in active mice, but they also displayed stronger satiety signaling during a negative energy-balance period.

To elucidate the gastrointestinal adaptations underlying the enhancement of the satiation, we conducted an acute 1-hour refeeding experiment post 12-hour daytime fasting. As expected, there were no differences in food intake (Figure 5D) or body weight gain (Figure 5E) between groups, even though active mice usually consume more during this time period (Figure S2H), anticipating that this results from the overcompensation seen in sedentary mice. Blood glucose increased after 1-hour of refeeding in both sedentary and active mice, but active mice had significantly lower glucose levels (Figure 5F). Next, we examined the gastrointestinal adaptations to fasting and refeeding in both sedentary and active mice. Whereas the wet weights of cecum and colon were not affected by refeeding or running, the wet weight of the stomach increased due to refeeding but not running, and the small intestinal wet weight tended to be greater in active mice (Figure S2D-G). Importantly, the length of small intestine and colon was greater due to running, in line with results shown in Figure 1F, and was not affected by feeding (Figure 5G). This highlights that small intestinal adaptations induced by running are not influenced by acute feeding-status. Through hematoxylin and eosin staining, we quantified villus height along the small intestine in fasted mice (Figure 5H). Villus height was similar between groups in each segment, although there may be an overall trend of higher villus if we do not correct for each segment (P=0.08 for the difference; Figure 5I). As expected, in both sedentary and active mice circulating GLP-1 and PYY levels increased whereas circulating ghrelin levels decreased in the refed-state (Figure 5J). Considering that active mice had lower circulating ghrelin and PYY during *ad-libitum* feeding (Figure 3C), this reveals that nutrient-stimulated peripheral signals affected by running are dependent of feeding-status.

### Increased physical activity induces molecular adaptations in the gut independently of feeding status, whereas it influences central nervous system adaptations in a feeding-status-dependent manner

Satiety is regulated by gut-to-brain communication.^39^ Thus, we investigated running-induced molecular adaptations in the gut-brain axis in response to 12-hour daytime fasting as well as acute (1-hour) and long-term (24-hour) refeeding. First, we measured the mRNA amounts of gut-peptides and their receptors in the proximal, middle, and distal small intestine. *Glp-1r* expression was upregulated in the proximal intestine, mainly at 1h- and 24-post-refeed. However, in the middle segment *Glp-1r* expression was downregulated in active mice independently of feeding-status, and further, *Glp-1r* expression trended to be downregulated in the refed- state in the distal small intestine (P=0.09 of the difference) (Figure 6A). Additionally, in the colon, *Glp-1r* expression was upregulated by running in the fasted and *ad-libitum*-fed state (Figure S2I). *The gene-code for pro-glucagon* (*Gcg*) and *Pyy* expression were upregulated along the small intestine. Although there was no overall effect of running on the expression of *Gcg*, it was downregulated in the middle and distal small intestine during refeeding (Figure 6B). *Pyy* expression was not regulated by running (Figure 6C). *Gpr65* expression, which is described to be important for nutrient sensing,^14,40,41^ was similar between sedentary and active mice (Figure 6D). Both *Cck* and its receptor, *Cckar*, were downregulated along the small intestine (Figure 6E-F). Running accentuated the downregulation of *Cckar*, in the middle small intestine, mainly during refeeding (Figure 6F). Overall, we observed that running regulates intestinal gene expression of *Glp-1r* and *Cckar* in a section-dependent manner, and *Glp-1r* expression was regulated independently of feeding-status.

**Figure 6.**
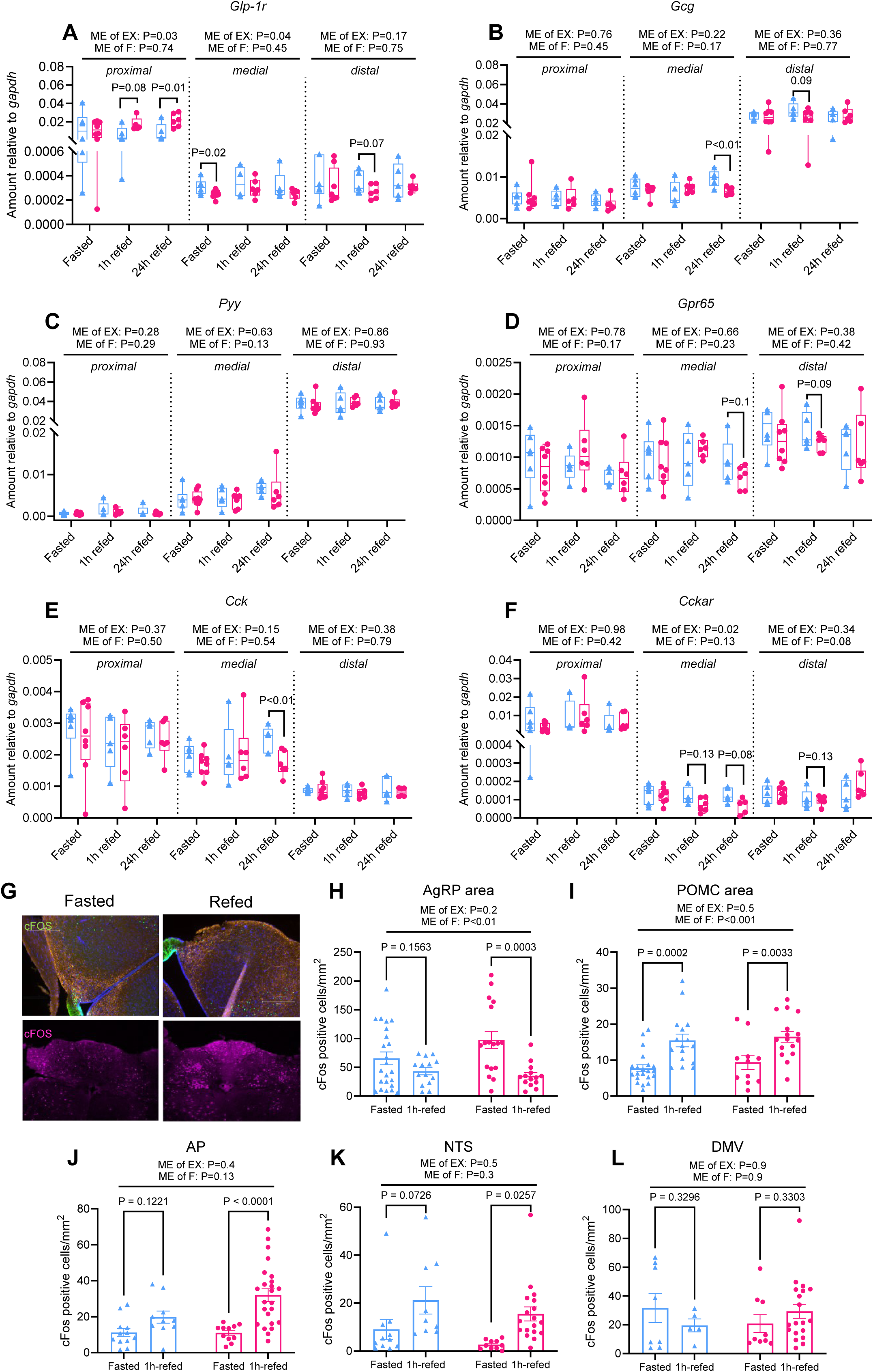
Training induced adaptations in gut-to-brain communication. Gene expression analysis through qPCR of proximal, middle and distal small intestine tissue in fasted, 1-hour refed and 24-hour refed mice; sedentary (blue) and active (pink). (A) *Glp-1r,* (B) *Gcg,* (C) *Pyy,* (D) *Gpr65,* (E) *Cck,* and (F) C*ckar* compared by two-way ANOVA with Bonferroni multiple comparison within each intestinal segment. (G) Immunofluorescence staining with cFOS antibody in the ARC (top) (CON-fasted *n*=3, CON-refed *n*=2, EX-fasted *n*=2, EX-refed *n*=4) and brainstem (bottom) (CON-fasted *n*=4, CON-refed *n*=4, EX-fasted *n*=3, EX-refed *n*=4). (H) cFOS positive cells/mm^2^ in the AgRP-area, (I) POMC-area of the ARC, and (J) in the AP, (K) NTS, (L) and DMV of the brainstem of fasted and 1-hour refed mice. Effect of feeding and exercise is compared by two-way ANOVA with Bonferroni multiple comparison. Data are shown as mean±SEM. P-values are depicted on the graphs. ME of EX is main effect of wheel running, ME of F is main effect of refeeding.

The *Glp1-r*, *Cckar* and *Gpr65* are expressed on vagal nerve innervating the gastrointestinal tract and signals are relayed through the nodose ganglia to the brainstem. Thus, we measured their expression in the nodose ganglia in response to 12-hour daytime fasting as well as 1-hour and 24-hour post-refeeding. Overall, *Glp1r* and *Cckar* expression was downregulated in active mice, and same trend was observed for *Gpr65* expression, with the difference in *Glp1r* expression being most pronounced during 1-hour post-refeeding (Figure S3D). In line with 24-hour post-refeeding results, there was no difference in GLP-1R+ positive neurons in the nodose ganglia when assessed by RNAscope in *ad-libitum*-fed mice (Figure S3E). Together, the data suggest that running induces nodose ganglia adaptations regardless of feeding-status. Signals from the nodose ganglia are relayed to brainstem. Therefore, we examined gene expression in the brainstem in *ad-libitum* feeding conditions during the pre-meal hunger (ZT12) and post-meal satiety (ZT0) phase to evaluate the effect of running in basal feeding state. Whereas running did not influence expression of *Glp-1r, Cckar, Cck* and *Gcg* in the pre-meal hunger state (ZT12) (Figure S3B), *Cckar* and *Gcg* expression were downregulated by running after overnight feeding (ZT0) (Figure S3C), which may be an adaptive satiation response to the observed gut adaptations. Since the feeding-related brain signals seems to be influenced by feeding-status and by physical activity levels, we finally assessed the acute neuronal activation in the brainstem and hypothalamus at fasted and 1h-refeed state (Figure 6G). In the hypothalamus, refeeding resulted in decreased neuronal activity in the AgRP-area and increased activity in the POMC-area (Figure 6H-I), as previously observed.^37^ The effect in neuronal activity of AgRP-area was more accentuated due to running, suggesting a stronger refeeding response. Further, in active mice there was a strong response to 1-hour refeeding in the AP and NTS of the brainstem, an effect that was not present in the sedentary mice (Figure 6J-K). In the DMV there was no clear effect of fasting and refeeding (Figure 6L). Overall, the data suggest that the dynamic regulation of feeding-related central signals is influenced by feeding-status and by physical activity levels.

## Discussion

Here we show that exercise-induced increase in food intake is associated with gastrointestinal tract adaptations that may represent underlying mechanism of the benefits of exercise on body weight maintenance.^42^ In particular, running induced an increase in small intestinal length, independently of GLP-2. Moreover, exercise enhanced sensitivity to gut-derived hormones, increased L-cell density and glucose-stimulated GLP-1 secretion and further differentially regulated intestinal *Glp-1r* expression. The increase in small intestine length was positively correlated with total running, and with total food intake in active but not sedentary mice, pointing to a key role of physical activity driving induced intestinal length. The intestine is the largest endocrine organ of the body and the gut-derived hormones have peripheral and central effects regulating energy homeostasis; therefore, the exercise-induced length may be of importance for its physiological functions.^43^ Previous work with obese mice showed that high-fat diet does not affect small intestine length,^44^ stating that intestine length is dependent on volume, not the calories of food ingested. Here, the small intestine lengthening induced by running was independent of acute changes in feeding-status, revealing that intestinal changes driven by increased energy expenditure may serve to chronically increase nutrient absorption capacity.

In humans, circulating GLP-1 increases in response to nutrient sensing and absorption^45^ and in response to an acute exercise bout.^46,47^ Moreover, habitual physical activity is associated with lower fasting and higher meal-induced GLP-1 secretion.^26^ In contrast, it has been demonstrated that people living with obesity have impaired pre- and post-meal GLP-1 secretion.^48–50^ In addition, *Glp-1* expression may be decreased in individuals living with obesity and in obese rodents.^51–53^ Here, running had no effect on circulating GLP-1 levels during *ad-libitum* feeding, but active mice had lower circulating ghrelin and lower circulating PYY levels compared to sedentary. Using the isolated perfused small intestine, we found that active mice had greater glucose-stimulated GLP-1 secretion compared to sedentary controls. Remarkably, running enhanced GLP-1 secretion independently of small intestine length (Figure S1C), in line with a greater number of GLP-1 positive cells per villus/crypt unit. This is, to our knowledge, the first evidence of direct exercise-induced regulation of GLP-1 production in the small intestine. In humans, acute exercise slows down gastric emptying,^54^ an effect possibly mediated by GLP-1.^29,55^ In the present study, increased physical activity trended to slow gastric emptying post-MMTT but not post-OGTT. Together these data suggest that physical activity induces changes in basal and meal-induced secretion of GLP-1, PYY, and ghrelin which may chronically improve appetite regulation.^20,30,56^

In accordance with this, we investigated sensitivity to gut-derived hormones related to food intake. Low doses of PYY and CCK suppressed food intake, and ghrelin induced food intake only in active mice, and this pattern was present during both the light and dark phases. Notably, exercise influenced feeding pattern by inducing food intake during the dark phase which resulted in increased total 24-hour food intake; however, notably, active mice ate less food during the light cycle than sedentary mice. Together, the data suggests that exercise impacts not only pre-meal hunger but also post-meal satiation and satiety. This is further supported by the present data showing stronger satiety signaling after daytime fasting and semaglutide-induced feeding suppression in active mice. In some human studies, self-reported hunger/satiety shows increased pre-meal hunger and post-meal satiety in people with high physical activity levels, and this was also consistent with plasma levels of total GLP-1 and acylated ghrelin.^30^ Further, it has been proposed that in people with high physical activity levels there is a stronger coupling between energy intake and expenditure; this conclusion was based on, among others, a high-energy preload study where physically active individuals were better at incorporating the calories of a calorie-dense preload meal in their total energy consumption so that their intake did not exceed energy needs.^30,57^ Which is coherent with the present results, where we observed a compensatory eating pattern in sedentary mice following daytime fasting resulting in additional body weight gain compared to the active mice. Overall, our study demonstrates that increased physical activity improves satiety signaling, especially during negative energy-balance periods, thereby contributing to the maintenance of a stable body weight.^58–60^

We assessed molecular changes in tissues related to appetite signaling. The *Glp1-r* and *Cckar* are highly expressed on gut-innervating enteric nerves and are found to be essential for sensitive appetite signaling by sensing nutrients and stretch in the gastrointestinal tract.^41,61–65^ In the present study *Glp-1r* expression was higher in the proximal small intestine and colon of active mice compared to sedentary mice and independent of feeding-status. In contrast, *Glp-1r* expression was downregulated by running in the middle part. GLP-1 is secreted in the same amount from the proximal and distal small intestine, and the intestinal feeding response is highly driven by proximally secreted GLP-1.^28,66^ We propose that the activity-induced increase in food intake drives an upregulation of *Glp-1r* expression in the proximal small intestine, which may be associated with a greater feeding response in active mice. Gut-derived signals stimulating the enteric nerves are relayed to the brainstem through the nodose ganglion. We propose here that running induces changes in intestinal gene expression which may results in adaptive responses in the nodose ganglia and brainstem. Indeed, *Glp-1r*, *Gpr65* and *Cckar* expression were downregulated in the nodose ganglia of active mice independently of feeding state, and further *Gcg, Cck and Cckar* expression were downregulated in the brainstem during the post-meal satiation phase, but not during pre-meal hunger phase in an *ad-libitum* feeding. To evaluate how feeding state could influence in the gut-derived hormones, we assessed them at fasted and refeed state in sedentary and active mice. There were no differences between the groups regarding circulating levels of fasting ghrelin nor post-meal PYY and GLP-1. In the hypothalamus, however, it seemed that induction of acute neuronal activity in AgRP-neuron was greater in the active mice during fasting compared to sedentary mice. In addition, there was a pattern of greater response to refeeding in the AgRP-area and brainstem of active mice compared to sedentary. Thus, even though circulating levels of appetite regulating hormones PYY, ghrelin and GLP-1 were similar between groups, their apparent ability to activate specific hypothalamic and brainstem neurons may be enhanced by running. However, more studies are needed to elucidate on the interaction between physical activity and acute neuronal activity in relation to appetite regulation.

In conclusion, increased energy expenditure resulting from running was accompanied by increased energy intake, which drives gastrointestinal adaptations modulating the intestinal responses to nutrients and enhancing the sensitivity to gut-derived peptides. Further, the number of GLP-1 positive L-cells was greater in active mice, suggesting a greater capacity for nutrient sensing and hormone secretion that may underlie the benefit of exercise on long-term appetite regulation and body weight maintenance.^67^ We propose that the morphological and molecular intestinal adaptations observed in active mice may be indicators of improved intestinal health and function. And that this results in more sensitive nutrient sensing and thus accurate the coupling between energy intake and expenditure, in turn driving the improvements on appetite regulation and body weight maintenance observed in individuals with high physical activity level.

### Limitations of the study

A limitation of the applied wheel running approach is the limited ability to control the volume, intensity and timing of the activity performed. However, this approach limits the risk of introducing a stress response as seen during treadmill or swimming exercises in mice.^68^ On the contrary, voluntary wheel running is rewarding and resembles the natural activity behavior in wild living mice.^68,69^ One outcome of the studies was neuronal activity in specific brain regions; thus, we chose a model of physical activity with low risk of stress responses that would be otherwise detected in these brain regions.

## STAR*METHODS

### Key resources table

| REAGENT or RESOURCE | SOURCE | IDENTIFIER |
| --- | --- | --- |
| **Antibodies** |  |  |
| GLP-1 rabbit-anti-mouse | Lab of Jens Juul Holst | 2135-8 |
| c-Fos rabbit-anti-mouse | Abcam | Cat# ab190289 |
| Donkey anti-Rabbit IgG (H+L) Highly Cross-Adsorbed Secondary Antibody, Alexa Fluor™ 647 | Thermo Fisher, DK | Cat # A-31573 |
| Donkey Anti-Rabbit IgG (H+L) Alexa Fluor® 555 antibody | Invitrogen | Cat # A-31572 |
| GLP-1 antibody | Lab of Jens Juul Holst | code no. 89390 |
| **Chemicals, peptides, and recombinant proteins** |  |  |
| Paraformaldehyde 4% | Thermo Fisher, DK | CAS: 30525-89-4 |
| TRIzol | Thermo Fisher, DK | CAS: 593-84-1 |
| SYBR Green Master Mix | Thermo Fisher, DK | Cat# 4309155 |
| Choloform | Merck | CAS: 67-66-3 |
| Peptide YY (PYY) | Novo Nordisk | NNC0165-1273 |
| Ghrelin | Biotechne | CAS: 258338-12-4 |
| Cholecystokinin (CCK) | Bachem | CAS: 25126-32-3 |
| GLP-1 (7-36) amide | Bachem | CAS: H-6795-GMP |
| Semaglutide | Novo Nordisk | NNC0113-0217 |
| I<sup>125</sup> GLP-1 | Novo Nordisk |  |
| I<sup>125</sup> PYY | Perkin Elmer | NEX2400 |
| PYY | Bachem | Cat# H-6045 |
| Hydrogen peroxide | Bio-techne | Cat# 322381 |
| Target retrieval | Bio-techne | Cat# 322000 |
| Protease plus | Bio-techne | Cat# 322331 |
| Label probes: Opal520, Cy3 and Cy5 | Perkin Elmer | Cat# NEL760001KT |
| Target probe: Mm-GLP1R-C3 | https://doi.org/10.1016/j.cmet.2021.05.002 | Cat# 418851-C3 |
| Target probe: Mm-Cckar-C1 | https://doi.org/10.1016/j.cmet.2021.05.002 | Cat#313751 |
| DAPI mounting medium VECTASHIELD | Vector Laboratories | CAT# H-1200 |
| 3-isobutyl-1-methylxanthine (IBMX) | Thermo Fischer, DK | CAT# 228420010 |
| Plasma-coated charcoal | Merck | Cat# 1.02186.0250 |
| **Critical commercial assays** |  |  |
| U-PLEX Obesity Combo 2 | MSD, Rockville USA | Cat# K15301K-1 |
| Acetaminophen L2K Assay | Sekure Chemistry | Cat# 506-10 |
| Multiplex Fluorescent Reagent Kit v2 | Advanced Cell Diagnostic | Cat# 323100 |
| **Experimental models: Organisms/strains** |  |  |
| Mouse: C57BL/6NRJ | Janvier | N/A |
| Mouse: C57BL/6NTac-Glp2rem5153Tac | In House (JJ Holst) | N/A |
| **Oligonucleotides** |  |  |
| See Table 1. | TAG Copenhagen<br>A/S | https://tagc.com/ |
| **Software and algorithms** |  |  |
| ImageJ | doi:10.1038/nmeth.2019 | https://imagej.nih.gov/ij/ |
| GraphPad Prism 10 | GraphPad Prism | https://www.graphpad.com/features |
| BDAS software |  | https://www.harvardapparatus.com/bdas-basic-data-acquisition-software.html |
| Wizard2, Automatic Gamma Counter | Perkin Elmer | N/A |
| EVOS™ M5000 Imaging System | Thermo Fisher, DK | N/A |
| **Other** |  |  |
| Chow diet | Safe Diets | SAFE D30 |

## EXPERIMENTAL MODEL AND SUBJECT DETAILS

### Ethical considerations

Animal studies were conducted with permission from the Danish National Committee for Animal Research (2020-15-0201-00599), from the local animal use committee (SUND, EMED, P24-183 and P24-252) and in accordance with the guidelines of Danish legislation governing animal experiments.

### Animals

Male C57BL/6NRJ mice (7 weeks old) were purchased from Janvier (Saint Berthevin Cedex, France). Global GLP-2 receptor knockout (6-8 weeks old) (GLP-2R^−/-^) mice were generated by Taconic, using CRISPR/Cas9-mediated gene editing, with Parent Designation: C57BL/6NTac-Glp2rem5153Tac. Animals were bred by heterozygote breeding and wild type (WT) littermates were used as controls. The mice were single-housed in a temperature (22±2°C) and humidity controlled (55%) room and kept in a 12:12h light/dark cycle with a preceding 30 min half-light period. The mice had *ad-libitum* access to standard chow (SAFE D30, Safe Diets, 3.389 kcal/g; 26% kcal energy from protein, 14% energy from fat, and 60% energy from carbohydrates) and drinking water.

## METHOD DETAILS

### Experimental setup

Body weight and food intake was measured daily, and through EchoMRI (EchoMRI^TM^ 4-in-1-500 Body Composition Analyzer, USA) scan body composition (fat -and lean mass) was measured once weekly. The mice were divided into matched groups based on these parameters: active (EX; voluntary wheel running) and sedentary (CON; blocked running wheel). Running wheels were opened after approximately one week of baseline measurements and mice were euthanized after 6 weeks of running. Statistical analysis information for each graph can be found in the figure legends. No statistical methods were applied to predetermine the sample size for experiments. All data are shown as mean ± SEM, and findings with *P ≤ 0.05, **P ≤ 0.01, ***P ≤ 0.001 were considered statistically significant.

**Study 1: Effect of voluntary wheel running on food intake, body weight, body composition and gastrointestinal tract.** Mice (7-weeks-old C57BL6/N, *n*=21 CON; *n*=21 EX) were in general kept on ad-libitum chow diet with daily measurement of body weight and food in the cage to calculate food intake. At 5-weeks of running mice were sacrificed to collect tissue: serum, small intestine, and colon.

**Study 2: Effect of fasting and refeeding on post-fasting food intake and body weight gain.** Mice (7-weeks-old C57BL6/N, *n*=14 CON; *n*=14 EX) were kept on ad-libitum chow diet with daily measurement of food in the cage to calculate food intake. At 4 weeks of running, body weight and food intake were measured, and food was removed from their cage just before the light cycle starts (06:00 AM). Body weight was measured, and food was re-introduced at 12-hours post-fasting (06:00 PM). Food intake and body weight were measured at 24-48- and 72-hours post-refeeding.

**Study 3: Effect of fasting and 1-hour refeeding on gene expression and neural activity**. Mice (7-weeks-old C57BL6/N, *n*=23 CON; *n*=27 EX) were kept on ad-libitum chow diet with daily measurement of food in the cage to calculate food intake. At 6 weeks, we performed terminal fasting and 1 h refeeding experiments. Body weight and food were measured at ZT0 whereafter cages were changed and food removed from cages and the animals kept fasting until ZT12. Mice were divided into two subgroups (euthanized in the fasted or refed state). Fasted mice were euthanized at ZT12 (*n*=12 CON-fasted, *n*=13 EX-fasted). Refed mice (*n*=11 CON-refed, *n*=14 EX-refed) were allowed to refeed for one hour before they were euthanized. Blood glucose was measured post killing and tissue was dissected and stored for future analysis: small intestine, colon, brain. Results from this study are combined from two cohorts of mice (*n*=50, 7-weeks-old C57BL6/N); they were exposed to the same handling and were euthanized in the same manner at the same age.

**Study 4: Effect of fasting refeeding; tissue collection.** Mice (7-weeks-old C57BL6/N n=42). Study 3 was repeated to supplement tissue collection for analysis. After 5-weeks of running, mice were euthanized in the following groups: CON-fasting *n*=6, CON-1-hour refed *n*=6, CON-24-hour refed *n=*6, EX-fasting *n=*8, EX-1-hour refed *n=*7, EX-24-hour refed *n=*8. Tissue collected: plasma, small intestine, colon, brain, and nodose ganglia.

**Study 5: Oral glucose tolerance test.** At 4 weeks of running mice from *study 4* underwent an oral glucose tolerance test. Mice were fasted from 06:00 AM (ZT0) to 12:00 PM (ZT6). At T = 0 mice received an oral gavage of a glucose (2 mg/kg BW) with acetaminophen (100 mg/kg) (Merck, CAS: 103-90-2). Blood glucose was measured in the tail vein by a hand-held glucometer at timepoint T = 0, 7.5, 15, 30, 60 and 90 min post oral gavage. At T = 0, 7.5 and 30-min post oral gavage 20 µl of blood was drawn from the tail vein into a capillary tube. Concentration of acetaminophen in the blood samples was used as an indicator of gastric emptying. Paracetamol was measured using a spectrophotometric method kit (Sekure Chemistry, Cat: 506-10).

**Study 6: Peptide hormone injections**. Peptide hormones related to feeding status were investigated in ad-libitum fed mice. The mice received intraperitoneal (i.p.) injection of peptide hormone PYY (0.004 µmol/kg BW) (Novo Nordisk, NNC0165-1273), CCK (3.8 nmol/kg BW) (Bachem, CAS: 25126-32-3), Ghrelin (0.3 µmol/kg BW) (bio-techne, CAS: 258338-12-4) to measure food induction/suppression and differences between groups. On the day of peptide administration food in all cages was measured 15 min before administration and removed from the cage. Body weight was measured to allow correct dosing and the food was introduced to the cage again shortly after the injection. Food was measured 1-, 2-, 4-, 12- and 24-hours post peptide administration. Body weight was measured at 12- and 24-hours post administration.

**Study 7: In situ small intestine perfusion.** Mice (7-weeks-old C57BL6/N, n=20) were kept on an ad-libitum chow diet with daily measurement of food in the cage to calculate food intake. At 5-weeks of running mice were used for in situ small intestine perfusion (n=8 CON; n=7 EX).

**Study 8: Mixed meal tolerance test.** At 4-weeks of running mice from *study 7* underwent an oral liquid mixed meal tolerance test (MMTT). Mice were fasted from 06:00 AM (ZT0) to 12:00 PM (ZT6). At T = 0 mice received an oral gavage of liquid mixed meal (0.2 mL; 2.4 kcal/mL) with acetaminophen (100 mg/kg BW). Blood glucose was measured from the tail vein by a hand-held glucometer at timepoint T = 0, 7.5, 15, 30, 60 and 90 min post oral gavage. To measure gastric emptying 20 µl of blood was drawn from the tail vein into a capillary tube at timepoint T = 0, 7.5 and 30-min post oral gavage. Concentration of acetaminophen in the blood samples was used as indicator of gastric emptying and was measured using a spectrophotometric method kit (Sekure Chemistry, Cat: 506-10).

**Study 9: Effect of running in GLP-2 KO**. Mice (6-8 weeks-old *n=*11 *glp2r*^−/-^*, n=*11 *glp2r^+/+^*) were kept on ad-libitum chow diet with daily measurement of food in the cage. At 5 weeks of running mice were sacrificed to collect: plasma, small intestine, and colon.

### Tissue and serum collection and processing

Mice were sacrificed by decapitation in the evening near lights off time. Feeding status at euthanasia is indicated in the figure legends. To ensure consistent results, the mice were euthanized in a randomized order, and further to minimize an effect of the circadian rhythm killing was sometimes performed over three days to ensure that the timing of euthanasia did not differ significantly. Blood was collected from trunk in and Eppendorf tube and quickly placed on ice. The small intestine, colon, brain, and nodose ganglia were surgically removed. Gastrointestinal tissue was weighed and measured before being divided into four sections: duodenum (proximal; first 2 cm after stomach), jejenum (medial; 2 cm approximately 15 cm from stomach), ileum (distal, last 2 cm before the cecum), and colon (last 2 cm before the rectum) (See figure 2). Depending on future analysis, tissue was either placed in 4% paraformaldehyde (PFA) or snap-frozen in liquid nitrogen.

### Serum analysis

As mentioned, whole-blood was collected post decapitation and plasma were saved at -80°C. Plasma samples were collected from *ad-libitum* fed mice (*study 1*), and from fasted versus 1-h refed mice (*study 3*) and analyzed for specific peptide hormones. U-PLEX Obesity Combo 2 (mouse) multiplex Assay (Meso Scale Discovery, Rockville, USA) was used following the manufacturer’s instruction to measure: total Ghrelin, total GLP-1, and total PYY. The kit also measures C-peptide, leptin, and insulin; however, these results will be published elsewhere.

### Gene expression analysis

Gene expression analysis was performed in small intestine, colon, and isolated brain stem samples. The tissue was surgically removed and snap-frozen with liquid nitrogen. Total RNA was isolated from tissues with phenol/chloroform extraction method. Briefly, tissue was homogenized with 1 ml of Trizol using Qiagen Tissuelyser Retsch MM300, 200µl chloroform was added to each sample and spun at 12,000 g for 10 min, 4 °C. After centrifugation, RNA was precipitated using isopropyl alcohol (1:1 reaction), washed with 75% ethanol and dissolved in Ultrapure RNase/DNase free water. RNA purity was measured using Nanodrop spectrophotometer (nanodrop one, Thermo Fischer Scientific, USA). RNA was converted into cDNA using cDNA Reverse Transcription Kit (Cat# 4368813, Life Technologies) following the instructions from the manufacturer. Quantitative PCR (qPCR) were performed with PowerUpTM SYBRTM Green Master Mix (Cat# A25780, ThermoFisher Scientific) and 300 nM of forward and reverse primers and using ViiA 7 Real Time PCR system (Applied Biosystems) for amplification. Relative quantification using ΔCt with a fold change normalization relative to the sedentary group was calculated. *Gadph* was used as housekeeping gene for all tissues.

**Table 1.**
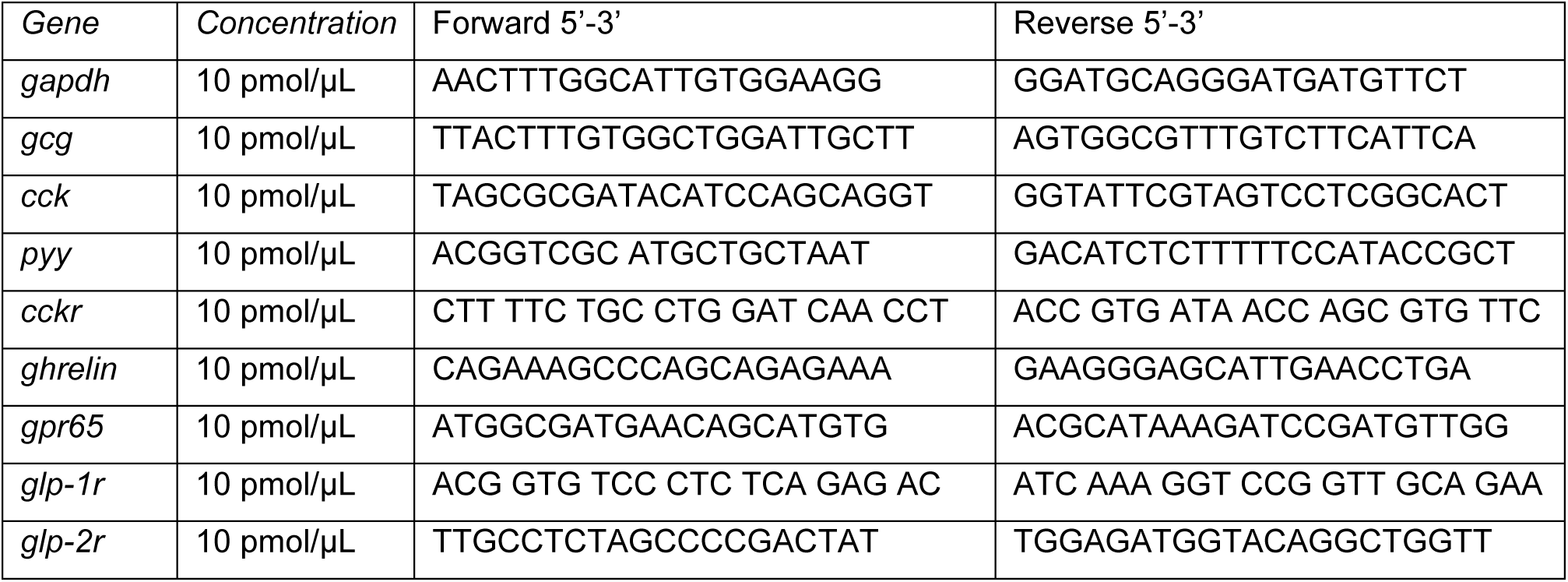
Primer sequences for qPCR.

### Hematoxylin and eosin staining with alcian blue

Intestinal tissue was surgically dissected, washed in PBS and placed in 4% PFA for 24 hours at RT. Samples were stored in 4% PFA at 4℃ and then embedded in paraffin and cut by In-Lab (Denmark, https://in-lab.dk/). HE-staining was used to quantify the height and depth of small intestinal villi and crypts. Histological images were acquired using EVOS5000 light microscopy. Villus height and crypt depth was measured manually with Fiji ImageJ. Two sections per intestinal segment from each mouse were analyzed. All villus and crypts were measured.

### Immunostaining and image analysis of small intestine and brain

The small intestine and brain were surgically removed, and the small intestine was divided in segments (proximal, middle and distal) before being placed in 4% (PFA) for 24 hours at RT. After 24 hours samples were moved to a 30% sucrose solution for 48-hours before being snap frozen in ice-cold isopentane and stored at -80 ℃. Intestinal tissue was cut in 16 µm thick sections and placed on glass slides, brains were cut in 20 µm thick slices and carefully placed in a liquid anti-freeze medium (30% phosphate buffer, 40% ethylene glycol and 30% glycerol) and stored at -20 ℃ until later use. Intestinal tissue samples stored on glass slides were used for IF staining targeting GLP-1 positive cells. IF were performed directly on the glass slides. Slides were blocked with a blocking solution (BS) (5% BSA, 5% normal donkey serum (NDS) in PBS) and dried with 100% ethanol before overnight incubation with primary GLP-1 AB (2135-8, 1:6000) at 4℃. Slides were washed with 1xPBS before second incubation with secondary AB (Alexa Fluor 647, 1:200) for 1 hour at RT in a dark room. After final incubation slides were washed and dried and added DAPI, a mounting media that stains cell nuclei, and hereafter stored at 4℃ until analysis.

Brain sections stored in anti-freeze media were washed with 1xPBS. Washing and incubation with AB was performed in wells, keeping brain sections wet at all times. Sections were placed in blocking solution (BS) followed by incubation in primary cFOS AB (ab190289, 1:500) overnight at 4℃ on a slow shaking table. Samples were washed and incubated with secondary AB (Alexa flour 647, 1:500) for 2 hours at RT in the dark. After the final wash, sections were mounted on glass slides and left to air dry in the dark. When slides were dry DAPI was added to each slide and covered with a coverslip. Cells positive for the target antigen were counted manually with Fiji ImageJ. Approx 6 brain and 3 intestine slides from each mouse were used for quantification.

### In situ hybridization of nodose ganglia

Nodose ganglia were dissected and post-fixed for 24-hours in PFA at RT following 48-hours incubation in 30% sucrose at 4°C. The ganglia were cut in 14 µm thick sections at -24°C and carefully places onto glass slides and stored at -20 ℃ until later use RNAscope. Multiplex Fluorescent Reagent Kit v2 (Advanced Cell Diagnostic, Cat# 323100) was used following the manufactures’ instructions. Sections on glass slides were dried at 60°C overnight. Hereafter they were pre-treated with hydrogen peroxide (Cat# 322381) and boiled in Target retrieval (Cat# 322000). Sections were then dehydrated in pure ethanol, a hydrophobic barrier was drawn around tissue sections (ImmEdge hydrophobic barrier pen, Vector Lab, H-4000), and the slides were then incubated for 15 min at 40°C in Protease Plus (Cat# 322331), followed by a 2 hour incubation in a HybEZ oven at 40°C with the target probes (Mm-Cckar-C1, Cat#313751; Mm-GLP1R-C3, Cat# 418851-C3). Signal was amplified with AMP1-3 and label probes (Opal520, Cy3 and Cy5; Perkin-Elmer, Cat# NEL760001KT). Lastly, sections were mounted using DAPI mounting medium (VECTASHIELD, Cat# H-1200, Vector Laboratories). Slides were imaged by a Zeiss ImagerM2 fluorescent microscope with 10x or 20x magnification or Leica TCA SP-8-X Confocal Microscope (Leica Microsystems) with 20x magnification. Images were processed using ImageJ and GIMPsoftware and neurons positive for RNAscope probes in nodose ganglia were quantified manually using ImageJ software from 2 different ganglia per mouse (n=4). Total number of neurons was assessed in each nodose ganglia using autofluorescence in FITC channels and colocalization of GLP1R and CCKaR mRNA was analyzed as percent relative to total number of neurons.

### In situ small intestine perfusion

*Ad-libitum fed mice* were anesthetized with an i.p. injection of a Ketamine/Xylazine mix (0.1 mL/20g) (Ketamine 90 mg/kg (Ketaminol Vet.; MSD Animal Health Madison, USA) and Xylazine (10 mg/kg (Rompun Vet.: Bayer Animal Health, Germany). The abdominal cavity was opened, and the small intestine was isolated by excising the surrounding organs: colon, stomach, and spleen, and the vasculature to the kidneys was ligated. A tube was inserted into the proximal small intestine just below the pyloric sphincter; the intestinal lumen was continuously perfused with heated (37°C) isotonic saline at flowrate 0.035 mL/min. A catheter was placed in the abdominal aorta perfusing the small intestine at 2.5 mL/min flowrate with a modified Krebs Ringer Bicarbonate buffer heated to 37°C and oxygenated with 95% O_2_ and 5% CO_2_. The venous effluent was collected by a catheter placed in vena portae. Hereafter, the mouse was euthanized by perforating the diaphragm. The intestine was perfused for 25 minutes before the initiation of the experimental protocol. Venous effluent samples were collected for 1 min periods. Samples were immediately placed on ice and stored at - 20°C until further analysis. Each protocol started with a 10 min baseline period of luminal saline infusion (0.035 mL/min) followed by intra-luminal glucose stimulation. The luminal stimuli were administered for 15 min at an initial bolus rate of 0.135 mL/min for 3-min (to replace previous solutions in the lumen) followed by 12-min at infusion rate 0.035 mL/min. Post glucose stimulus the lumen was infused with saline to allow wash out bolus rates mentioned above. Intra-vascular KCl infusion at 50 mmol/L (final concentration) was used as a positive control in the end of each experiment.

#### Perfusion system

We used a *single-pass* perfusion system (Uniper UP-100, Hugo Electronics-Harvard apparatus, March-Hugstetten, Germany) which heats the perfusion buffer to 37°C). The perfusion buffer was a modified Krebs-Ringer bicarbonate buffer containing 0.1% (w/v) bovine serum albumin (BSA) (Merck KGaA, Darmstadt, Germany), 5% (w/v) dextran T-70 to balance osmolarity (Pharmacosmos, Denmark), 3.5 mmol/L glucose and 5 mmol/L of fumarate, glutamate and pyruvate, and 10 µmol/L IBMX. The buffer was constantly gassed with 95% 0_2_/5% CO_2_ to maximally increase pO_2_ and maintain pH around 7.4. Arterial as well as venous perfusate samples were analyzed regularly for pH and partial pressures of oxygen and carbon dioxide (Radiometer Acid-Base laboratory). The intestinal perfusion method is described in detail elsewhere.^70^

#### Biochemical measurements of perfusion effluent

Total amidated GLP-1 and total PYY concentrations in venous effluents from intestinal perfusions were measured by in-house-developed radioimmunoassay (RIA). Total GLP-1 (7-36amide) was measured with an in-house-developed RIA, based on a C-terminally directed antiserum specific for the amidated GLP-1 form (code no. 89390).^71^ The standard was synthetic GLP-1 7–36NH_2_ (cat. no. H-6795-GMP, 4081700, Bachem, Frechen, Germany), and the tracer was monoiodinated ^125^I-labeled GLP-1 (7–36NH2) (a gift from Novo Nordisk A/S, Bagsværd, Denmark). Total PYY was measured with a porcine antiserum (cat. no T-4093, Bachem, Germany); it should be noted that porcine and murine PYY share amino acid sequence.^72^ The antibody used recognizes PYY1/3– 36 and PYY1/3–34 equally. The standard was synthetic rat/mouse/porcine PYY (cat. no. H-6045, Bachem, Germany), and the tracer was ^125^I-labeled porcine PYY (cat. no. NEX2400). Free and bound peptides were separated with plasma-coated charcoal (cat. no. 1.02186.0250, Merck, Germany). Antigen-bound peptide radioactivity was counted by a gamma counter (Wizard2, Automatic Gamma Counter, PerkinElmer, Denmark), and radioactivity was translated into hormone levels by interpolation on a standard curve with known concentrations increasing from 0 pmol/L to 320 pmol/L an adequate range to detect all samples in this study.

### Data presentation and statistical analysis

The graphs were made using GraphPad prism 9 (GraphPad, La Jolla, USA). All data are presented as mean±SEM. Statistical analyses were made using GraphPad and SPFF. Statistical analysis employed is noted in figure legends. P-values<0.05 was considered significant. P-values are depicted on the graphs.

## ACKNOWLEGDEMENTS

Anne Jørgensen, Lene Foged and Ida Holm are acknowledged for their technical assistance. This study was supported by a research grant from the Novo Nordic Foundation (grant ID 0059436) and the Centre for Physical Activity Research (CFAS), which is an independent research center at Rigshospitalet and is supported by Trygfonden (grants ID 101390, ID 20045, ID 125132, and ID 177225). P.S is supported by Lundbeck Foundation grant (grant ID R380-2021-1300).

## AUTHOR CONTRIBUTION

Conceptualization, C.B-L., J.J.H, B.K.P, and P.S.; Methodology, C.B-L., J.J.H. and P.S; Formal analysis, C.B-L. and P.S.; Investigation, C.B-L., J.V.U., K.D.G., J.L., H.L.K and P.S; Visualization, C.B-L.; Writing – original draft, C.B-L.; Writing – review & editing, C.B-L., J.J.H., B.K.P., and P.S.; Supervision, C.B-L., J.J.H, B.K.P. and P.S.

## RESOURCE AVAILABILITY

### Lead Contact

Further information requests regarding resources and reagents should be directed to and will be answered by the Lead Contact, Paula Sanchis

### Materials Availability

This study did not generate new unique materials.

### Data and Code Availability

Raw data is available upon request to lead contact Paula Sanchis. This study dod not generate any codes.

## SUPPLEMANENT INFORMATION TITLES AND LEGENDS

Document 1. Figures 1-6

Document S1. Supplementary Figures 1-3.

## SUPPLEMNTARY FIGURE LEGENDS

**Supplementary Figure 1. Food intake following peptide injection.** (A) PYY secretion from the perfused small intestine at baseline and during luminal glucose stimulation, and (B) total PYY secretion at baseline and glucose-stimulated PYY secretion compared by two-way ANOVA. (C) Total glucose-stimulated GLP-1 secretion from the perfused small intestine normalized based on small intestine length. (D) Accumulated food intake following injection of PYY, and (E) following CCK in different doses compared by one-way ANOVA. (F) Fold change of food intake following two doses of ghrelin injection related to vehicle injection. (G) Hourly food intake following GLP injection (15 nmol/kg BW). Quantification of cFos positive cells in the (H) ARC, (I) AP, and (J) NTS following injection of either, saline, PYY, or Ghrelin. Gene expression analysis of *Gcg, Pyy* and *Cck* in the saline injection group of (K) proximal, (L) middle, and (M) distal small intestine compared by t-test. Data are shown as mean±SEM. P-values are depicted on the graphs.

**Supplementary Figure 2. Effect of calorie-restriction on food intake and gastrointestinal adaptations.** (A) Color scheme of figure. (B) Difference in food intake between baseline and 12-, 24-, 48-, and 72-hours post semaglutide injection was compared by Bonferroni multiple comparison. Relative difference to baseline food intake and P-values are depicted on the graph. (C) Body weight change (%) post semaglutide injection compared by Bonferroni multiple comparison. Wet weight of (D) small intestine, (E) colon, (F) cecum, and (G) stomach of fasted or 1-hour refed mice compared by two-way ANOVA and Bonferroni multiple comparison. (H) Food intake during 1-hour refeeding (ZT12-ZT13) post 12-hour daytime fasting, and 1-hour food intake (ZT12-ZT13) during *ad-libitum* feeding conditions compared by t-test. (I) Gene expression of *Gcg, Glp-1r,* and *Glp-2r* in the colon of fasted, 1-hour refed, or 24-hour refed mice compared by mixed model with multiple comparison. Data are shown as mean±SEM. P-values are depicted on the graphs. ME of EX is main effect of wheel running, ME of F is main effect of refeeding.

**Supplementary Figure 3. Running-induced regulatory expression in vagal signaling pathways.** (A) Gene expression analysis of *Gcg, Glp-1r,* and *Glp-2r* in colon tissue of sedentary and active mice during *ad-libitum* feeding at ZT12 compared by t-test. (B) Gene expression analysis of *Glp-1r, Gcg, Gpr65, Cck,* and *Cckar,* in brainstem tissue during ZT12, and (C) gene expression of *Glp-1r, Gcg, Cck,* and *Cckar* in the brainstem at ZT0. (D) Gene expression analysis through qPCR of G*lp-1r, Gpr65,* and *Cckar* in nodose ganglia compared by two-way ANOVA with Bonferroni multiple comparison. (E) RNAscope analysis of GLP-1R-positive neurons in the nodose ganglia compared by t-test (CON *n*=3, EX *n*=2). Data are shown as mean±SEM. P-values are depicted on the graphs.

## REFERENCES

1. Hall KD, Kahan S. Maintenance of Lost Weight and Long-Term Management of Obesity. Med Clin North Am. 2018;102(1):183–197. doi:10.1016/j.mcna.2017.08.012

2. Catenacci VA, Ogden LG, Stuht J, et al. Physical Activity Patterns in the National Weight Control Registry. Obesity (Silver Spring). 2008;16(1):153–161. doi:10.1038/oby.2007.6

3. Kolnes KJ, Petersen MH, Lien-Iversen T, Højlund K, Jensen J. Effect of Exercise Training on Fat Loss—Energetic Perspectives and the Role of Improved Adipose Tissue Function and Body Fat Distribution. Front Physiol. 2021;12(September):1–14. doi:10.3389/fphys.2021.737709

4. Donnelly JE, Honas JJ, Smith BK, et al. Aerobic exercise alone results in clinically significant weight-loss for men and women: Midwest exercise trial 2. Obesity. 2013;21(3):219–228. doi:10.1002/oby.20145

5. Franz MJ, VanWormer JJ, Crain AL, et al. Weight-Loss Outcomes: A Systematic Review and Meta-Analysis of Weight-Loss Clinical Trials with a Minimum 1-Year Follow-Up. J Am Diet Assoc. 2007;107(10):1755–1767. doi:10.1016/j.jada.2007.07.017

6. Blundell JE, Gibbons C, Caudwell P, Finlayson G, Hopkins M. Appetite control and energy balance: Impact of exercise. Obesity Reviews. 2015;16(S1):67–76. doi:10.1111/obr.12257

7. Mayer J, Roy P, Mitra KP. Relation between Caloric Intake, Body Weight, and Physical Work. Am J Clin Nutr. 1956;4(2):169–175. doi:10.1093/ajcn/4.2.169

8. Martins C, Kulseng B, King NA, Holst JJ, Blundell JE. The Effects of Exercise-Induced Weight-loss on Appetite-Related Peptides and Motivation to Eat. J Clin Endocrinol Metab. 2010;95(4):1609–1616. doi:10.1210/jc.2009-2082

9. Beaulieu K, Hopkins M, Blundell J, Finlayson G. Does Habitual Physical Activity Increase the Sensitivity of the Appetite Control System? A Systematic Review. Sports medicine (Auckland). 2016;46(12):1897–1919. doi:10.1007/s40279-016-0518-9

10. Clemmensen C, Müller TD, Woods SC, Berthoud HR, Seeley RJ, Tschöp MH. Gut-Brain Cross-Talk in Metabolic Control. Cell. 2017;168(5):758–774. doi:10.1016/j.cell.2017.01.025

11. Sun EWL, Martin AM, Young RL, Keating DJ. The regulation of peripheral metabolism by gut-derived hormones. Front Endocrinol (Lausanne). 2019;10(JAN):1–11. doi:10.3389/fendo.2018.00754

12. Wachsmuth HR, Weninger SN, Duca FA. Role of the gut–brain axis in energy and glucose metabolism. Exp Mol Med. 2022;54(4):377–392. doi:10.1038/s12276-021-00677-w

13. Grau-Bove C, Gonzalez-Quilen C, Cantini G, et al. GLP1 Exerts Paracrine Activity in the Intestinal Lumen of Human Colon. Int J Mol Sci. 2022;23(7):3523. doi:10.3390/ijms23073523

14. Zanchi D, Depoorter A, Egloff L, et al. The impact of gut hormones on the neural circuit of appetite and satiety: A systematic review. Neurosci Biobehav Rev. 2017;80:457–475. doi:10.1016/j.neubiorev.2017.06.013

15. Abbott C, Monteiro M, Small C, et al. The inhibitory effects of peripheral administration of peptide YY3-36 and glucagon-like peptide-1 on food intake are attenuated by ablation of the vagal-brainstem-hypothalamic pathway. Brain Res. 2005;1044(1):127–131. doi:10.1016/j.brainres.2005.03.011

16. Modvig IM, Christiansen CB, Rehfeld JF, Holst JJ, Veedfald S. CCK-1 and CCK-2 receptor agonism do not stimulate GLP-1 and neurotensin secretion in the isolated perfused rat small intestine or GLP-1 and PYY secretion in the rat colon. Physiol Rep. 2020;8(2):e14352-n/a. doi:10.14814/phy2.14352

17. Martinez de Morentin PB, Gonzalez JA, Dowsett GKC, et al. A brainstem to hypothalamic arcuate nucleus GABAergic circuit drives feeding. Current biology. 2024;34(8):1646–1656.e4. doi:10.1016/j.cub.2024.02.074

18. Alhadeff AL. Monitoring In Vivo Neural Activity to Understand Gut–Brain Signaling. Endocrinology (Philadelphia). 2021;162(5):1. doi:10.1210/endocr/bqab029

19. Drazen DL, Vahl TP, D’Alessio DA, Seeley RJ, Woods SC. Effects of a Fixed Meal Pattern on Ghrelin Secretion: Evidence for a Learned Response Independent of Nutrient Status. Endocrinology (Philadelphia). 2006;147(1):23–30. doi:10.1210/en.2005-0973

20. Janus C, Vistisen D, Amadid H, et al. Habitual physical activity is associated with lower fasting and greater glucose-induced GLP-1 response in men. Endocr Connect. 2019;8(12):1607–1617. doi:10.1530/EC-19-0408

21. Bailey DM, Davies B, Castell LM, Newsholme EA, Calam J. Physical exercise and normobaric hypoxia: independent modulators of peripheral cholecystokinin metabolism in man. Journal of applied physiology (1985). 2001;90(1):105–113. doi:10.1152/jappl.2001.90.1.105

22. Zouhal H, Sellami M, Saeidi A, et al. Effect of physical exercise and training on gastrointestinal hormones in populations with different weight statuses. Nutr Rev. 2019;77(7):455–477. doi:10.1093/nutrit/nuz005

23. Ouerghi N, Feki M, Bragazzi NL, et al. Ghrelin Response to Acute and Chronic Exercise: Insights and Implications from a Systematic Review of the Literature. Sports medicine (Auckland). 2021;51(11):2389–2410. doi:10.1007/s40279-021-01518-6

24. Anderson KC, Zieff G, Paterson C, Stoner L, Weltman A, Allen JD. The effect of acute exercise on pre-prandial ghrelin levels in healthy adults: A systematic review and meta-analysis. Peptides (New Yorkm NY : 1980). 2021;145:170625–170625. doi:10.1016/j.peptides.2021.170625

25. Mailhac A, Pedersen L, Pottegård A, et al. Semaglutide (Ozempic®) Use in Denmark 2018 Through 2023 ‒ User Trends and off-Label Prescribing for Weight-loss. Clin Epidemiol. 2024;16:307–318. doi:10.2147/CLEP.S456170

26. Lundgren JR, Janus C, Jensen SBK, et al. Healthy Weight-loss Maintenance with Exercise, Liraglutide, or Both Combined. New England Journal of Medicine. 2021;384(18):1719–1730. doi:10.1056/nejmoa2028198

27. Litvak DA, Hellmich MR, Evers BM, Banker NA, Townsend CM. Glucagon-Like Peptide 2 Is a Potent Growth Factor for Small Intestine and Colon. Journal of Gastrointestinal Surgery. 1998;2(2):146–150. doi:10.1016/S1091-255X(98)80005-X

28. Svendsen B, Pedersen J, Albrechtsen NJW, et al. An analysis of cosecretion and coexpression of gut hormones from male rat proximal and distal small intestine. Endocrinology (United States). 2015;156(3):847–857. doi:10.1210/en.2014-1710

29. Holst JJ, Gribble F, Horowitz M, Rayner CK. Roles of the Gut in Glucose Homeostasis. Diabetes Care. 2016;39(6):884–892. doi:10.2337/dc16-0351

30. Long SJ, Hart K, Morgan LM. The ability of habitual exercise to influence appetite and food intake in response to high- and low-energy preloads in man. British Journal of Nutrition. 2002;87(5):517–523. doi:10.1079/bjn2002560

31. Larraufie P, Roberts GP, McGavigan AK, et al. Important Role of the GLP-1 Axis for Glucose Homeostasis after Bariatric Surgery. Cell reports (Cambridge). 2019;26(6):1399–1408.e6. doi:10.1016/j.celrep.2019.01.047

32. Chen XY, Chen L, Yang W, Xie AM. GLP-1 Suppresses Feeding Behaviors and Modulates Neuronal Electrophysiological Properties in Multiple Brain Regions. Front Mol Neurosci. 2021;14:793004–793004. doi:10.3389/fnmol.2021.793004

33. Drucker DJ. Mechanisms of Action and Therapeutic Application of Glucagon-like Peptide-1. Cell Metab. 2018;27(4):740–756. doi:10.1016/j.cmet.2018.03.001

34. Pradhan G, Samson SL, Sun Y. Ghrelin: much more than a hunger hormone. Curr Opin Clin Nutr Metab Care. 2013;16(6):619–624. doi:10.1097/MCO.0b013e328365b9be

35. Small CJ, Herzogk H, Cohen MA, et al. Gut hormone PYY 3-36 physiologically inhibits food intake. Nature. 2002;418(August):728–730. https://idp.nature.com/authorize/casa?redirect_uri= https://www.nature.com/articles/nature00887&casa_token=3AU8AVRKV3YAAAAA:QFBFVbrr987mRQljNGqfNxoX_wDM413EBMsL8jsUfZYuXoJc7t9fyeNetRtZA0RvgtLmx3F1XAne86jXZA

36. Jones ES, Nunn N, Chambers AP, Østergaard S, Wulff BS, Luckman SM. Modified Peptide YY Molecule Attenuates the Activity of NPY/AgRP Neurons and Reduces Food Intake in Male Mice. Endocrinology. 2019;160(11):2737–2747. doi:10.1210/en.2019-00100

37. Wu Q, Lemus MB, Stark R, et al. The Temporal Pattern of cfos Activation in Hypothalamic, Cortical, and Brainstem Nuclei in Response to Fasting and Refeeding in Male Mice. Endocrinology (Philadelphia). 2014;155(3):840–853. doi:10.1210/en.2013-1831

38. Zheng Z, Zong Y, Ma Y, Tian Y. Glucagon-like peptide-1 receptor: mechanisms and advances in therapy. Signal Transduct Target Ther. 2024;(June). doi:10.1038/s41392-024-01931-z

39. Barakat GM, Ramadan W, Assi G, Khoury NB El. Satiety: a gut–brain–relationship. The journal of physiological sciences. 2024;74(1):11–11. doi:10.1186/s12576-024-00904-9

40. Bai L, Mesgarzadeh S, Ramesh KS, et al. Genetic Identification of Vagal Sensory Neurons That Control Feeding. Cell. 2019;179(5):1129–1143.e23. doi:10.1016/j.cell.2019.10.031

41. Borgmann D, Ciglieri E, Biglari N, et al. Gut-brain communication by distinct sensory neurons differently controls feeding and glucose metabolism. Cell Metab. 2021;33(7):1466–1482.e7. doi:10.1016/j.cmet.2021.05.002

42. Shook RP, Hand GA, Drenowatz C, et al. Low levels of physical activity are associated with dysregulation of energy intake and fat mass gain over 1 year. Am J Clin Nutr. 2015;102(6):1332–1338. doi:10.3945/ajcn.115.115360

43. Bany Bakar R, Reimann F, Gribble FM. The intestine as an endocrine organ and the role of gut hormones in metabolic regulation. Nat Rev Gastroenterol Hepatol. 2023;20(12):784–796. doi:10.1038/s41575-023-00830-y

44. Xie Y, Ding F, Di W, et al. Impact of a high-fat diet on intestinal stem cells and epithelial barrier function in middle-aged female mice. Mol Med Rep. 2020;21(3):1133–1144. doi:10.3892/mmr.2020.10932

45. Holst JJ. The physiology of glucagon-like peptide 1. Physiol Rev. 2007;87(4):1409–1439. doi:10.1152/physrev.00034.2006

46. Åkerström T, Stolpe MN, Widmer R, et al. Endurance Training Improves GLP-1 Sensitivity and Glucose Tolerance in Overweight Women. J Endocr Soc. 2022;6(9):1–8. doi:10.1210/jendso/bvac111

47. Ellingsgaard H, Seelig E, Timper K, et al. GLP-1 secretion is regulated by IL-6 signalling: a randomised, placebo-controlled study. Diabetologia. 2020;63(2):362–373. doi:10.1007/s00125-019-05045-y

48. Skibicka KP. The central GLP-1: implications for food and drug reward. Front Neurosci. 2013;7:181–181. doi:10.3389/fnins.2013.00181

49. Williams DL. Neural integration of satiation and food reward: Role of GLP-1 and orexin pathways. Physiol Behav. 2014;136:194–199. doi:10.1016/j.physbeh.2014.03.013

50. Berthoud HR. Metabolic and hedonic drives in the neural control of appetite: who is the boss? Curr Opin Neurobiol. 2011;21(6):888–896. doi:10.1016/j.conb.2011.09.004

51. Matikainen N, Bogl LH, Hakkarainen A, et al. GLP-1 Responses Are Heritable and Blunted in Acquired Obesity With High Liver Fat and Insulin Resistance. Diabetes Care. 2014;37(1):242–251. doi:10.2337/dc13-1283

52. Faerch K, Torekov SS, Vistisen D, et al. GLP-1 Response to Oral Glucose Is Reduced in Prediabetes, Screen-Detected Type 2 Diabetes, and Obesity and Influenced by Sex: The ADDITION-PRO Study. Diabetes (New York, NY). 2015;64(7):2513–2525. doi:10.2337/db14-1751

53. Dusaulcy R, Handgraaf S, Skarupelova S, et al. Functional and Molecular Adaptations of Enteroendocrine L-Cells in Male Obese Mice Are Associated With Preservation of Pancreatic α-Cell Function and Prevention of Hyperglycemia. Endocrinology (Philadelphia). 2016;157(10):3832–3843. doi:10.1210/en.2016-1433

54. Lehrskov LL, Christensen RH, Wedell-Neergaard AS, et al. Effects of Exercise Training and IL-6 Receptor Blockade on Gastric Emptying and GLP-1 Secretion in Obese Humans: Secondary Analyses From a Double Blind Randomized Clinical Trial. Front Physiol. 2019;10:1249–1249. doi:10.3389/fphys.2019.01249

55. Holst JJ, Madsbad S. Mechanisms of surgical control of type 2 diabetes: GLP-1 is key factor. Surgery for obesity and related diseases. 2016;12(6):1236–1242. doi:10.1016/j.soard.2016.02.033

56. Grau-Bove C, Gonzalez-Quilen C, Cantini G, et al. GLP1 Exerts Paracrine Activity in the Intestinal Lumen of Human Colon. Int J Mol Sci. 2022;23(7):3523. doi:10.3390/ijms23073523

57. Martins C, Truby H, Morgan LM. Short-term appetite control in response to a 6-week exercise programme in sedentary volunteers. British Journal of Nutrition. 2007;98(4):834–842. doi:10.1017/S000711450774922X

58. Beaulieu K, Hopkins M, Blundell J, Finlayson G. Does Habitual Physical Activity Increase the Sensitivity of the Appetite Control System? A Systematic Review. Sports medicine (Auckland). 2016;46(12):1897–1919. doi:10.1007/s40279-016-0518-9

59. Gibbons C, Blundell J. Appetite regulation and physical activity – an energy balance perspective. Hamdan Medical Journal. 2015;8(1):33. doi:10.7707/hmj.431

60. Quist JS, Klein AB, Færch K, et al. Effects of acute exercise and exercise training on plasma GDF15 concentrations and associations with appetite and cardiometabolic health in individuals with overweight or obesity – A secondary analysis of a randomized controlled trial. Appetite. 2023;182:106423–106423. doi:10.1016/j.appet.2022.106423

61. Williams EK, Chang RB, Strochlic DE, Umans BD, Lowell BB, Liberles SD. Sensory Neurons that Detect Stretch and Nutrients in the Digestive System. Cell. 2016;166(1):209–221. doi:10.1016/j.cell.2016.05.011

62. Vasiliki Vana, Michelle K. Lærke, Jens F. Rehfeld, et al. Vagal afferent cholecystokinin receptor activation is required for glucagon-like peptide-1–induced satiation. Published online 2022.

63. Winzenried ET, Neyens DM, Calkins R, Appleyard SM. CCK-expressing neurons in the NTS are directly activated by CCK-sensitive C-type vagal afferents. Am J Physiol Regul Integr Comp Physiol. 2025;328(1):R121. doi:10.1152/ajpregu.00280.2023

64. Sayegh AI, Ritter RC. CCK-A receptor activation induces Fos expression in myenteric neurons of rat small intestine. Regul Pept. 2000;88(1):75–81. doi:10.1016/S0167-0115(99)00124-X

65. Grunddal K V, Jensen EP, Ørskov C, et al. Expression Profile of the GLP-1 Receptor in the Gastrointestinal Tract and Pancreas in Adult Female Mice. Endocrinology (Philadelphia). 2022;163(1):1. doi:10.1210/endocr/bqab216

66. Borg MJ, Bound M, Grivell J, et al. Comparative effects of proximal and distal small intestinal administration of metformin on plasma glucose and glucagon-like peptide-1, and gastric emptying after oral glucose, in type 2 diabetes. Diabetes Obes Metab. 2019;21(3):640–647. doi:10.1111/dom.13567

67. Shah M, Vella A. Effects of GLP-1 on appetite and weight. Rev Endocr Metab Disord. 2014;15(3):181–187. doi:10.1007/s11154-014-9289-5

68. Skovbjerg G, Fritzen AM, Svendsen CSA, et al. Atlas of exercise-induced brain activation in mice. Molecular metabolism (Germany). 2024;82:101907–101907. doi:10.1016/j.molmet.2024.101907

69. Meijer JH, Robbers Y. Wheel running in the wild. Proceedings of the Royal Society B: Biological Sciences. 2014;281(1786). doi:10.1098/rspb.2014.0210

70. Jepsen SL, Grunddal K, Albrechtsen NJW, et al. Paracrine crosstalk between intestinal L- and D-cells controls secretion of glucagon-like peptide-1 in mice. Am J Physiol Endocrinol Metab. 2019;317(6):E1081–E1093. doi:10.1152/ajpendo.00239.2019

71. Ørskov C, Rabenhøj L, Wettergren A, Kofod H, Holst JJ. Tissue and plasma concentrations of amidated and glycine-extended glucagon-like peptide I in humans. Diabetes. 1994;43(4):535–539. doi:10.2337/diab.43.4.535

72. Toräng S, Veedfald S, Rosenkilde MM, Hartmann B, Holst JJ. The anorexic hormone peptide yy3-36 is rapidly metabolized to inactive peptide yy3-34 in vivo. Physiol Rep. 2015;3(7):1–8. doi:10.14814/phy2.12455

